# Natural rubber reduces herbivory and alters the microbiome below ground

**DOI:** 10.1101/2022.10.10.511194

**Authors:** Laura Böttner, Antonino Malacrinò, Christian Schulze Gronover, Nicole van Deenen, Boje Müller, Shuqing Xu, Jonathan Gershenzon, Dirk Prüfer, Meret Huber

**Author notes:** **Correspondence:** Meret Huber, Dirk Prüfer.

## Abstract

- Laticifers are hypothesized to mediate both plant-herbivore and plant-microbe interactions. However, there is little evidence for the dual function of these secretory structures.
- We investigated whether the major constituent of natural rubber, *cis*-1,4-polyisoprene, a phylogenetically widespread and economically important latex polymer, alters plant resistance and the root microbiome of the Russian dandelion (*Taraxacum koksaghyz)* under attack of a root herbivore, the larva of the May cockchafer (*Melolontha melolontha)*.
- Rubber-depleted transgenic plants lost more shoot and root biomass upon herbivory than normal rubber content near-isogenic lines. *M. melolontha* preferred to feed on artificial diet supplemented with rubber-depleted rather than normal rubber content latex. Likewise, adding purified *cis*-1,4-polyisoprene in ecologically relevant concentrations to diet deterred larval feeding and reduced larval weight gain. Metagenomics and metabarcoding revealed that abolishing biosynthesis of natural rubber alters the structure but not the diversity of the rhizosphere and root microbiota in a herbivore-dependent manner. Roots from rubber-depleted plants, however, did not exhibit a higher pathogen load compared to normal rubber content roots.
- Taken together, our data demonstrate that natural rubber biosynthesis reduces herbivory and alters the plant microbiota in a herbivore-dependent manner, which highlights the role of plant specialized metabolites and secretory structures in shaping multitrophic interactions.

## Introduction

Tissue integrity and wound sealing is vital for all organisms as wounds constrain tissue functioning and facilitate the entry of pathogens (Savatin *et al*., 2014). In vertebrates, wounding often involves the disruption of blood vessels, and consequently the draining blood carries defensive cells, antibodies and coagulants to the site of wounding. Similar wound sealing and defensive processes are evident in laticifers, specialized cells found in about 10% of all flowering plants. Laticifers store their often milky cytoplasm, called latex, under pressure (Metcalfe, 1967; Farrell *et al*., 1991; Thomas M. Lewinsohn, 1991). Upon damage, latex exudes and can thereby deter or impair the growth of herbivores, or even kill them outright (Dussourd, 1995; Konno *et al*., 2004; Bont *et al*., 2017). Consequently, latex may reduce plant damage and improve plant fitness under herbivory (Agrawal & Konno, 2009; Konno, 2011; Huber *et al*., 2016b,a; Salomé Abarca *et al*., 2019; Castelblanque *et al*., 2021; Gracz-Bernaciak *et al*., 2021).

Apart from defense against herbivores, laticifers are hypothesized to mediate plant-microbe interactions (Agrawal & Konno, 2009). Experimental evidence for this notion is however scarce. Mutants of *Euphorbia lathyris* that lack laticifers showed enhanced resistance towards the fungal leaf pathogen *Botrytis cinerea*, possibly due to the presence of microbial growth stimulating factors in laticifers (Castelblanque *et al*., 2021). Whether latex mediates plant-microbe interactions upon exuding through wounds is, however, unclear. Considering that laticifers are among the most common defensive reservoirs in plants (Thomas M. Lewinsohn, 1991), this knowledge gap significantly constrains our understanding of plant constitutive defenses against microbes.

Latex typically contains high concentrations of specialized metabolites (Sessa *et al*., 2000; Agrawal & Konno, 2009; Huber *et al*., 2015; Salomé-Abarca *et al*., 2021) that are known to serve as defenses against herbivores (Steppuhn *et al*., 2004; Züst *et al*., 2012; Kerwin *et al*., 2015; Huber *et al*., 2016b; Li *et al*., 2018). These and other specialized metabolites are also increasingly recognized as mediators of plant-microbe interactions (Schütz *et al*., 2021). For instance, specialized metabolites show growth-promoting or growth-inhibiting activities towards microbes (Pang *et al*., 2021), and transgenic plants deficient in specialized metabolites can be more susceptible to pathogens (Tierens *et al*., 2001; Bednarek *et al*., 2009; Koprivova *et al*., 2019; Chen *et al*., 2020). Furthermore, abolishing the production of specialized metabolites, such as coumarins, benzoxazinoids and triterpene derivatives, through genetic manipulations has revealed their role in altering the composition and function of the root and rhizosphere microbiome (Stringlis *et al*., 2018; Hu *et al*., 2018; Cotton *et al*., 2019; Huang *et al*., 2019; Voges *et al*., 2019; Jacoby *et al*., 2021).

Plant-microbe interactions however do not occur in isolation, but may be modulated by the presence of herbivores. For instance, pathogens often enter through wounds made by feeding herbivores (Friedli & Bacher, 2001; Savatin *et al*., 2014), and microbes may use herbivores as vectors to enter plants (Willsey *et al*., 2017). Furthermore, signalling networks during plant-microbe and plant-herbivore interactions are partially shared and interact (Koornneef & Pieterse, 2008; Robert-Seilaniantz *et al*., 2011; Hilleary & Gilroy, 2018). Despite the close interaction of plants, herbivores and microbes, the impact of plant specialized metabolites on the microbiome in the presence or absence of herbivores is rarely assessed (Hu *et al*., 2018).

A latex metabolite that has frequently been hypothesized to mediate both plant-microbe and plant-herbivore interactions is natural rubber, which consists of over 90% *cis*-1,4-polyisoprene accompanied by other species-dependent compounds. Natural rubber is found in almost 10% of all latex-bearing plants, albeit in different quantities, molecular weights and structures with different branching (Mooibroek & Cornish, 2000; Agrawal & Konno, 2009). Natural rubber is among the economically most important plant polymers (Mooibroek & Cornish, 2000), and has likely convergently evolved implicating important ecological functions (Metcalfe, 1967; Agrawal & Konno, 2009). The most commonly postulated function involves herbivore defense since natural rubber is sticky and thus may entrap entire insects or glue their mouthparts (Dussourd & Eisner, 1987; Dussourd, 1995). Direct evidence that natural rubber is involved in herbivore defense is however scarce.

Apart from herbivore defense, natural rubber may also contribute to wound sealing, thereby preventing the entry of pathogenic microorganisms. Wound sealing may be particularly important below-ground, as the soil harbors a rich microbiota including pathogens (Tringe *et al*., 2005; Fierer & Jackson, 2006). When coagulated on a solid growth medium, c*is*-1,4-polyisoprene blocked the entry of pathogenic bacteria and delayed the entry of fungi *in vitro* (Salomé-Abarca *et al*., 2021). Whether natural rubber alters the plant microbiota and restricts the entry of pathogenic microorganisms *in planta* is, however, unclear.

The Russian dandelion (*Taraxacum koksaghyz*, Asteraceae) - an obligate outcrossing diploid - accumulates particularly high quantities of high-molecular weight natural rubber in its laticifers, especially in roots with levels of 2-12% of root dry weight (Ulmann, 1951). The concentration of *cis*-polyisoprene in the roots is reduced by almost 90% after transgenic knock-down of the expression of a *cis*-prenyltransferase-like subunit (CPTL1), a protein that is required for rubber chain elongation in the laticifers (Niephaus *et al*., 2019). *T. koksaghyz*, native to Kazakhstan and the western part of Xinjiang (China), is increasingly cultivated as a rubber crop in Europe, where it is attacked by the soil-dwelling herbivorous larvae of the May cockchafer, *Melolontha melolontha* (Scarabeidae, Coleoptera) (Härdtl, 1953). *M. melolontha*, native to Europe, is polyphagous like many Scarabeidae larvae that occur in the native range of *T. koksaghyz* (Gninenko, 1998; Malysh *et al*., 2002; Jackson & Klein, 2006), although its preferred host plants are dandelions (Hauss & Schütte, 1976).

Here we address the hypotheses that *cis*-1,4-polyisoprene (1) improves resistance of *T. koksaghyz* to *M. melolontha* herbivory, (2) alters the plant microbiome and (3) reduces pathogen load under *M. melolontha* attack. Using transgenic rubber-deficient lines, chemical supplementation and behavioral assays, we demonstrate that *cis*-1,4-polyisoprene helps protect *T. koksaghyz* from *M. melolontha* herbivory. Furthermore, amplicon sequencing and shotgun metagenomics revealed that the biosynthesis of this polyisoprene may alter the root and rhizosphere microbiome, but not the pathogen load in a herbivore-dependent manner.

## Material & Methods

### Plant material and growth conditions

*Taraxacum koksaghyz* and *Daucus carota ssp. sativus* (Nantaise2/Jubila, Kiepenkerl, Bruno Nebelung GmbH) were cultivated in a glasshouse with temperature 20–25°C, 20 klx light intensity (high pressure sodium lamp, HPS 600 Watts, Greenbud, enhanced yellow and red spectrum) with a 16h photoperiod. Two independently transformed *Tk*CPTL1-RNAi lines (RNAi = “rubber-depleted”) and their respective rubber-bearing near isogenic lines (NILs = “normal rubber content”) were used (Niephaus *et al*., 2019). For details on lines and cultivation see Supporting Information Methods S1-2.

### Insect material

*Melolontha melolontha* larvae were collected from meadows in Switzerland (46.692442, 9.410272; Urmein, Viamala region), and Germany (49.936182, 9.279764; Mespelbrunn, Spessart region). Experiments were performed with the third instar larvae (L3). Insects were reared individually in 180ml plastic beakers filled with a mix of potting soil and grated carrots in 24h darkness at 12-14°C. For experiments, larvae were starved for 72 h in the dark at room temperature. All *ex planta* experiments were performed in the dark to avoid light disturbance.

### Data analysis

Data analysis was performed with R Version 4.1.2 (R Core Team, 2020) and visualized with ggplot2 (Wickham, 2016). Details to the procedures are described in the individual sections.

### Resistance of rubber-depleted *Tk*CPTL1-RNAi and rubber-bearing NIL plants under *M. melolontha* herbivory

To test whether natural rubber benefits *T. koksaghyz* under root herbivory, two rubber-depleted (*Tk*CPTL1-RNAi-A and -B) lines and their respective normal rubber content NIL controls (NIL-A and -B) (plant_genotype=rubber-depleted or normal rubber content) were subjected to *M. melolontha* herbivory. Plants were cultivated for 10 weeks, a time point at which NIL plants accumulate natural rubber to 2-3% per root dry weight (Niephaus *et al*., 2019). Half of the replicates were then infested with one pre-weighed *M. melolontha* larva. After 15 days, plant roots and shoots were harvested and weighed, and larval weight gain was measured. Experiments with lines from the A and B genetic backgrounds were performed at different time points. Since the A lines suffered high infestation of white flies and thrips during the experiment, they were excluded from the analysis. Plants that spontaneously flowered, heavily wilted due to loss of the main root during *larval* feeding, or whose larva died during the experiment were also excluded from analysis. Shoot and root fresh weights were analyzed with linear models using the formula *‘biomass∼plant_genotype*herbivory_treatment’*. Pairwise comparisons between treatments within each plant genotype were adjusted using the FDR method with the package emmeans (Lenth, 2022) using the formula ‘pairwise∼herbivory_treatment|plant_genotype’. Difference in larval weight gain between normal rubber content and rubber-depleted plants was analyzed with a Wilcoxon signed-rank test.

### Choice experiment with carrot seedlings supplemented with latex

To investigate whether natural rubber alters *M. melolontha* feeding preference, we recorded the choice of *M. melolontha* larvae between carrot seedlings coated with latex of either *Tk*CPTL1-RNAi or NIL plants. Roots of five-week-old carrot seedlings were covered entirely with latex of freshly cut 10-week-old *Tk*CPTL1-RNAi or NIL *T. koksaghyz* roots of the A and B genetic backgrounds. Seedlings painted with *Tk*CPTL1-RNAi and NIL latex were pairwise arranged on opposite sides of 180ml beakers filled with vermiculite (n=23 A-pairs; n=28 B-pairs). One larva was then placed in the center of each beaker. Larval position (*Tk*CPTL1-RNAi side, NIL side, inactive=no choice) was recorded every hour until six to seven hours after start of the experiment. Experiments on A and B plants were performed at two separate time points. Choice data was analyzed for each line and time point separately using a binomial test.

### Choice experiment with carrot seedling supplemented with *cis*-1,4-polyisoprene

To test whether *cis*-1,4-polyisoprene deters *M. melolontha* feeding, we assessed the choice of the larvae between carrot seedlings supplemented with ecologically relevant concentrations of purified *cis*-1,4-polyisoprene (purification protocol in Methods S3) or a solvent control. Soil adhering to eight-week-old carrot seedlings was washed off, and seedlings arranged in pairs of homogeneously looking root sizes (n=50). Carrot roots were dipped three times either in 1% w/v purified *cis*-1,4-polyisoprene dissolved in chloroform or chloroform (solvent control), resulting in approximately 1.1% *cis*-1,4-polyisoprene based on root fresh weight, similar to concentrations of natural rubber in roots of wildtype *T. koksaghyz* (0.2-1.2% of root fresh weight (Ulmann, 1951)). Carrot seedlings of each treatment were placed on opposite sides of 180ml beakers filled with vermiculite. One *M. melolontha* larva was placed in the center of each beaker, and larval position (*cis*-1,4-polyisoprene side, solvent control side, inactive=no choice) was recorded after 1h, 2h, 3h, 4h, 6h. Choice data was analyzed for each line and time point separately using a binomial test.

### Non-choice experiment with artificial diet supplemented with *cis*-1,4-polyisoprene

To investigate whether natural rubber alters larval growth, we fed *M. melolontha* larvae for five consecutive days with artificial diet supplemented with isolated *cis*-1,4-polyisoprene (Supporting Information Methods S3) or solvent as a control. Artificial diet cubes of 400 mg (Huber *et al*., 2016b) were supplemented with either 1.2ml of a 1.1% w/v *cis*-1,4-polyisoprene solution in chloroform or chloroform as a solvent control, resulting in 3% *cis*-1,4-polyisoprene in the rubber-supplemented cubes. Cubes were incubated under a fume hood for 1h to allow chloroform to fully evaporate before the feeding assays. Pre-weighed larvae were allowed to feed solitarily on a diet cube for 24h inside a 180ml plastic beaker covered with a moist tissue (n=26 per treatment). The procedure was repeated with the same individuals for five consecutive days. Larvae that did not feed throughout the experiment were excluded from the analysis. Diet consumption was measured daily and data was analyzed by fitting a linear mixed effect model with the package *lme4* using the formula *‘consumption∼supplementation_treatment*(1*|*day)’*. Total larval weight gain after five days, and total weight gain per total consumed diet were analyzed with Wilcoxon signed-rank tests.

### Role of triterpenes in *T. koksaghyz* resistance and *M. melolontha* behavior

As roots of rubber-depleted *Tk*CPTL1-RNAi plants accumulate about three fold higher pentacyclic triterpene levels compared to NIL plants (Niephaus *et al*., 2019), we also investigated whether triterpenes alter *T. koksaghyz* performance under *M. melolontha* herbivory using two independent triterpene-reduced *TkOSC*-RNAi lines (oxidosqualene cyclase knock-down; *TkOSC*-RNAi-L2 and -L3) and their respective NILs (van Deenen *et al*., 2019). The *TkOSC*-RNAi lines displayed 73-80% reduced pentacyclic triterpene content, but normal *cis*-1,4-polyisoprene levels (van Deenen *et al*., 2019). In addition, we assessed whether lupeol, a *T. koksaghyz* triterpene, alters *M. melolontha* growth when added to artificial diet in physiologically relevant amounts (Unland *et al*., 2018; Pütter *et al*., 2019). Details on the experimental setup and statistical analysis can be found in Supporting Information Methods S4.

### Microbial colonization the of root-soil continuum upon herbivory

To investigate whether natural rubber biosynthesis alters the microbial colonization of roots and the rhizosphere upon damage, we compared root and rhizosphere microbiomes of rubber-depleted *Tk*CPTL1-RNAi -A and normal rubber content NIL-A plants (‘plant genotype’) under control conditions, *M. melolontha* herbivory and mechanical wounding (‘treatment’) using amplicon sequencing and shotgun metagenomics. Mechanical wounding was included to impose a uniform damage treatment that is unaffected by *Tk*CPTL1-RNAi silencing. All plants were propagated in soil originating from an agricultural field in which *T. koksaghyz* had been cultivated for several years (for details see Supporting Information Methods S5). Plants were either non-infested (NIL n=10, *Tk*CPTL1-RNAi n=10), infested with one pre-weighed *M. melolontha* larva (NIL n=9, *Tk*CPTL1-RNAi n=10), or wounded using a metal stick pricking into the pots 10 times every third day (NIL n=9, R *Tk*CPTL1-RNAi n=9). At 14 days after infestation, plants were harvested, root and shoot fresh weight determined and biomass data analyzed as described above. For assessing the microbiome, we chose a subset of six plants per treatment that showed visual signs of damage upon herbivory or wounding. To obtain the rhizosphere fraction, roots were washed in three consecutive steps with 30ml sterile water similar to (Hu *et al*., 2018), and washing fractions were combined. Roots were immediately frozen in liquid nitrogen, stored at – 20°C, subsequently lyophilized and ground to a fine powder. Details can be found in the Supporting Information Methods S6.

### DNA extraction, library preparation and sequencing

DNA was isolated from the lyophilized rhizosphere fraction and root powder using the DNeasy Powersoil Pro kit (Qiagen, Hilden, Germany) according to the manufacturer’s instructions. To account for possible contaminations, a negative control was included where the root powder was replaced by the same amount of ultrapure water. Extracted DNA was quantified and checked for purity using a spectrophotometer (Pearl 3435, Implen, Munich, Germany) and submitted to Novogene (Beijing, China) for library preparation and sequencing.

#### Shotgun metagenomics

The genomic DNA was randomly fragmented by sonication, then DNA fragments were end-polished, A-tailed, and ligated with full-length adapters for Illumina sequencing, followed by PCR amplification with P5 and indexed P7 oligos. The PCR products as the final construction of the libraries were purified using the AMPure XP reagents. Then libraries were checked for size distribution by an Agilent 2100 Bioanalyzer (Agilent Technologies, CA, USA), and quantified by qPCR. Libraries were then sequenced on an Illumina NovaSeq 6000 instrument (Illumina, CA, USA) on a S4 150PE flow cell.

#### Amplicon sequencing

DNA extracted from the rhizosphere and roots was processed to generate an amplicon library targeting the 16S (bacteria) and ITS2 (fungi) regions of the rRNA. Briefly, after samples passed QC, libraries were built by amplifying the 16S rRNA gene (515F and 806R primers) or the ITS2 rRNA gene (ITS3-2024F and ITS4-2409R primers). Purified PCR products (Beckman and Coulter AMPure XP) were used for a second PCR to ligate Illumina adapters. Libraries were purified again, pooled at equimolar ratio, and sequenced on Illumina NovaSeq 6000 instrument (Illumina, CA, USA) on a SP 250PE flow cell.

### Bioinformatics data processing and analysis

#### Shotgun metagenomics

Raw data was processed with TrimGalore v0.6.6 (Krueger, 2020) to remove adapters and low-quality reads, and reads obtained from the host plant were discarded through Bowtie2 v2.4.4 (Langmead & Salzberg, 2012) using the *T. koksaghyz* genome (Lin *et al*., 2018). Reads were merged into contigs using *MegaHit* v1.2.9 (Li *et al*., 2015) and functional annotation was performed using Prokka v1.14.6 (Seemann, 2014). Raw reads were mapped against the output from Prokka using Bowtie2 and Samtools (Danecek *et al*., 2021) obtaining a count matrix of gene frequency for each sample.

#### Amplicon sequencing

De-multiplexed forward and reverse reads were merged using the PEAR 0.9.1 algorithm with default parameters (Zhang *et al*., 2014). Data QC, OTU (operational taxonomic unit) clustering and chimera removal were carried out using VSEARCH 2.14.2 (Rognes *et al*., 2016). Taxonomy was assigned to each OTU using VSEARCH by querying the SILVA database (v. 138) (Quast *et al*., 2013). Singletons and OTUs coming from amplification of plastidial DNA were discarded from the downstream analyses.

### Microbiome data analysis

#### Diversity analysis

Shannon and Simpson diversity indices were calculated for each sample using the package *phyloseq* (McMurdie & Holmes, 2013). Then a generalized linear model was fit using Bayesian Hamiltonian Markov chain Monte Carlo in the package *rstanarm* (Goodrich *et al*., 2020), using the formula ‘*metric∼plant_genotype*treatment’* where the term *metric* represented either Shannon or Simpson index. Weakly informative normally distributed priors were used for both the intercept (mean=0, scale=2.5) and coefficients (mean=0, scale=2.5), with autoscaling turned on, running four chains with 2000 iterations each and discarding the first 1000 as burn-in, for a total of 4000 observations. Comparisons between groups were performed using the function hypothesis of the *brms* R package (Bürkner, 2017). The equivalent of a two-tailed *P*-value (pMCMC) was estimated by calculating and doubling the frequency at which any of the 4000 observations disagreed with (had opposite sign to) the posterior estimate.

#### Multivariate analysis

In order to perform the structure analysis, the OTU counts were normalized using DESeq2 (Love *et al*., 2014), and then fit to a Bayesian Generalized Linear Multivariate Multilevel Model in the *brms* package using the formula ‘*∼plant_genotype*treatment* ‘, running four chains with 2000 iterations each and discarding the first 1000 as warm-up, for a total of 4000 observations. Comparisons between groups were performed using the function hypothesis of the *brms* package (Bürkner, 2017); the equivalent of a two-tailed *P*-value (pMCMC) was estimated as reported above.

#### Differential taxa

Each plant genotype (NIL-A and *Tk*CPTL1-RNAi-A) and each treatment (herbivory or wounding) was contrasted towards the respective control group using *DESeq2*, keeping the OTUs that resulted being differentially abundant within each treatment/control combination (FDR-adjusted *P*<0.05). At the same time, a similar set of data was generated, but the counts were randomized within the OTU table, while keeping the number of reads within each sample constant. This enabled to test whether the number of differentially abundant OTUs in each group was different from random using a chi-squared test. Similarly, we tested whether treatment (within each of the plant genotypes) influences the relative abundance of OTUs grouped for each microbial genus. To this end, data were aggregated at the ‘Genus’ taxonomic level, and filtered to remove those genera that represent less than 1% of the community within each group. Then, the influence of treatment on the relative abundance of each bacterial genus for each plant genotype was tested separately by fitting a linear model using the *lme4* package. We used a chi-squared test to test the effect of plant genotype on the number of unique OTUs of each treatment (wounding, herbivory) compared to the respective control group for each compartment (root, rhizosphere) separately.

#### Magnitude of change

Each plant genotype (NIL-A and *Tk*CPTL1-RNAi-A) and each treatment (herbivory or wounding) was contrasted towards the respective control group using *DESeq2*. Then abs(log2FC) values were then compared by fitting a linear mixed effect model with the package *lme4* using the formula ‘group * (1|geneID)’ where *group* indicates the four combinations of genotypes (NIL-A and *Tk*CPTL1-RNAi-A) and treatments (herbivory or wounding). Pairwise comparisons were then tested using the package *emmeans*.

#### Gene content analysis

The gene count table was normalized using *DESeq2* and then fit to a Bayesian Generalized Linear Multivariate Multilevel Model as described above. Also, genes differentially abundant between each treatment (herbivory or wounding) / control combination, within each plant genotype, were identified using *DESeq2* and their number was tested compared to a random null model as described above.

## Results

### *Cis*-1,4-polyisoprene benefits *T. koksaghyz* under *M. melolontha* attack

To test whether the biosynthesis of *cis*-1,4-polyisoprene, the dominant constituent of natural rubber, benefits plant performance upon root herbivory, we subjected plants of transgenic *cis*-1,4-polyisoprene-depleted *Tk*CPTL1-RNAi lines (“rubber-depleted”) and the corresponding NIL lines (“normal rubber-content”) to *M. melolontha* herbivory. Rubber-depleted *Tk*CPTL1-RNAi plants suffered stronger reduction in shoot growth under *M. melolontha* herbivory compared to the rubber-bearing NIL plants (interaction *plant_genotype*herbivory, P*=0.03, linear model, Fig. 1a). In NIL plants, *M. melolontha* herbivory did not affect shoot growth (*P*=0.9, FDR corrected pairwise contrast, Fig. 1a), whereas in *Tk*CPTL1-RNAi plants, *M. melolontha* reduced shoot biomass by 33% (*P*=0.003, FDR corrected pairwise contrast, Fig. 1a). A similar pattern was observed for root biomass: in NIL plants, *M. melolontha* herbivory did not affect root biomass (*P*=0.6, FDR corrected pairwise contrast, Fig. 1b), whereas in *Tk*CPTL1-RNA plants, *M. melolontha* reduced root biomass by 34% (*P*=0.02, FDR corrected pairwise contrast, Fig. 1b). *M. melolontha* larvae gained approximately two-fold more weight on the *Tk*CPTL1-RNAi than on the NIL plants, but these differences were statistically not significant due to the large within-group variation (*P*≥0.13, Wilcoxon test, Fig. 1c). Originally, RNAi and control NIL lines of two different genetic backgrounds, A and B, were tested. Only data for the B background, however, are presented here. Plant resistance in the A background could not be analyzed as growth was dwarfed due to high levels of white fly and thrips infestation. However, in a follow-up experiment, the A background RNAi plants exhibited a similar albeit only marginally significant resistance pattern compared to the corresponding A NIL lines as that described above for the B background, (Supporting information, Fig. S5).

**Figure 1.**
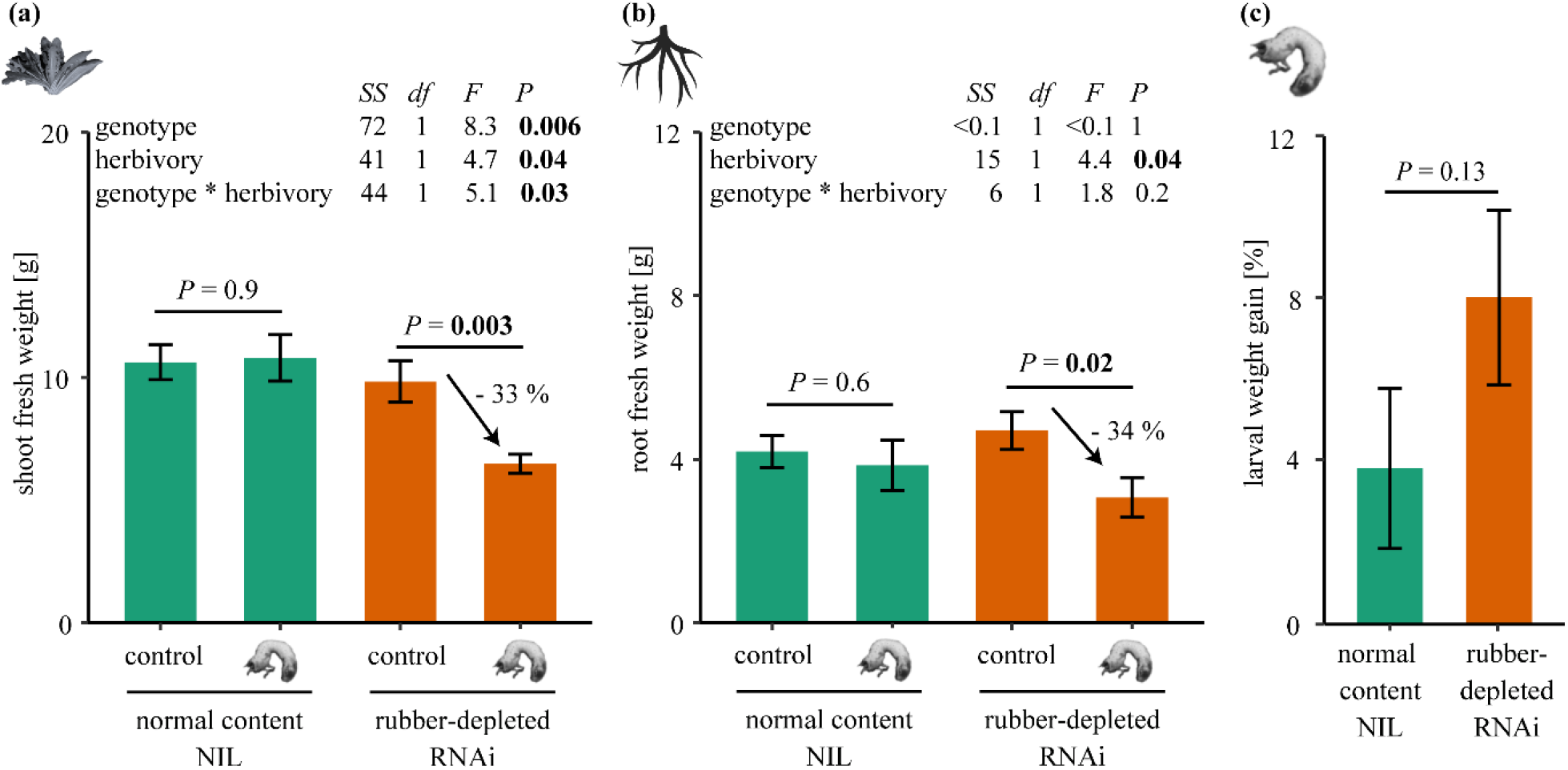
Silencing the biosynthesis of the major natural rubber metabolite, *cis*-1,4-polyisoprene, reduces the performance of *Taraxacum koksaghyz* under *Melolontha melolontha* herbivory. **(a)** Shoot and **(b)** root fresh weight accumulation of *T. koksaghyz* after 15 days of *M. melolontha* herbivory in rubber-depleted *Tk*CPTL1-RNAi lines and rubber-bearing NIL plants (“plant_genotype”). *P*-values of linear models are shown on top of the panels. *P*-values of pairwise comparisons were adjusted using the FDR method. **(c)** *M. melolontha* weight gain on *Tk*CPTL1-RNAi and NIL plants. *P*-values of Wilcoxon signed-rank test is shown. SS=sum squares, df=degrees of freedom, N=11-18. Error bars=standard error of means.

To corroborate the role of natural rubber in herbivore defense, we performed a series of *in planta* and *ex planta* experiments. First, we assessed *M. melolontha* feeding preference between carrot seedlings coated with latex of *Tk*CPTL1-RNAi or their respective NIL plants. For both the A and B lines, larvae preferred to feed on carrot seedlings painted with latex of *Tk*CPTL1-RNAi plants (*P*<0.05, binomial tests) with approximately 75% of the larvae choosing the rubber-deficient plants (Fig. 2a). Larval choice showed an expected temporal pattern, with a gradual increase of the number of larvae on the rubber-depleted side, followed by a neutral distribution of the larvae (Supporting Information Fig. S1).

**Figure 2.**
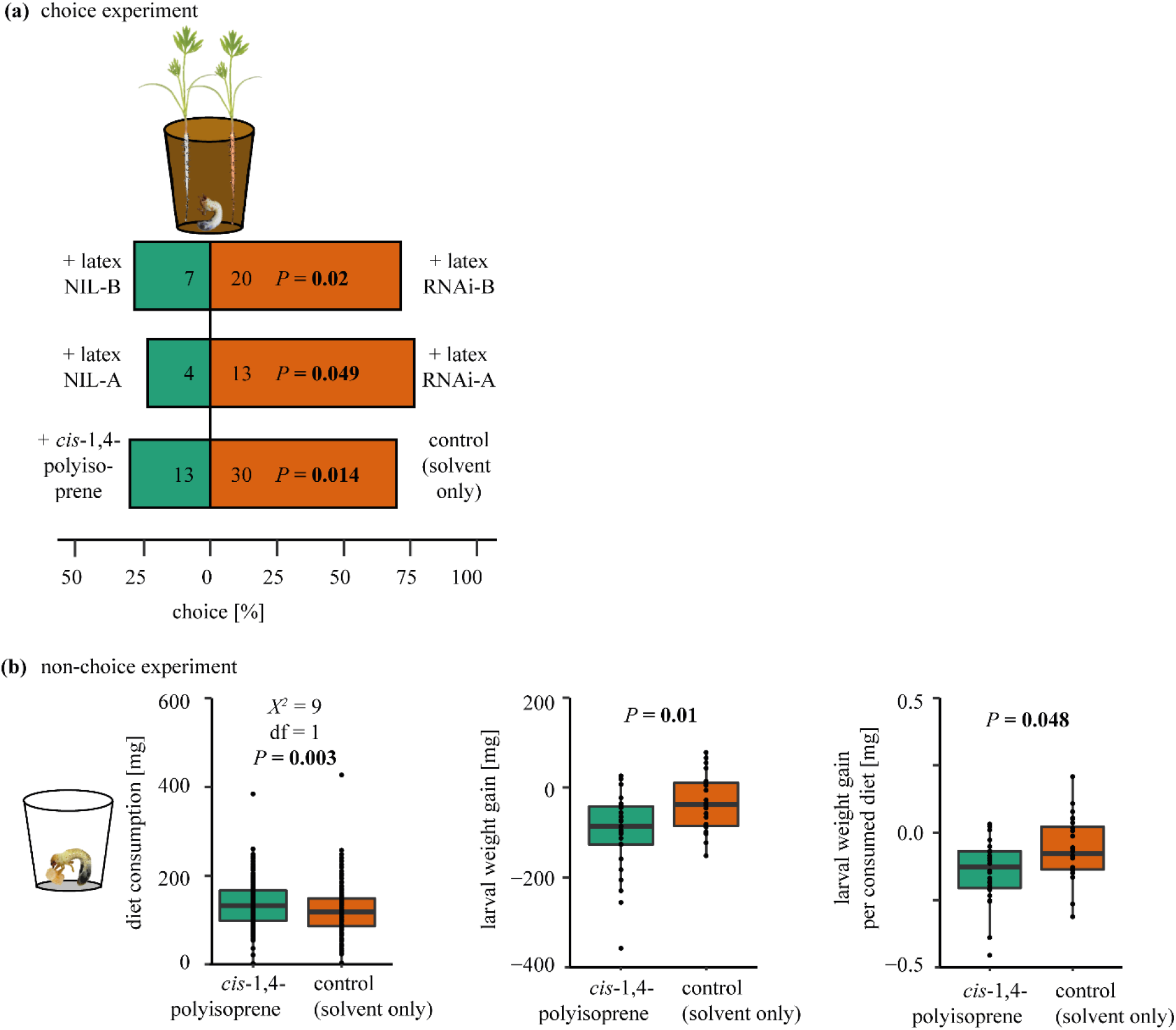
Ecologically relevant concentrations of *cis*-1,4-polyisoprene deter *Melolontha melolontha* feeding and reduce food quality. **(a)** Choice of *M. melolontha* larvae between carrot seedlings supplemented with latex of rubber-depleted *Tk*CPTL1-RNAi plants and normal rubber content NIL plants, or between seedlings supplemented with 1.1% *cis*-1,4-polyisoprene and a solvent control. *P*-values of binomial tests are shown. The number of active larvae is indicated inside the bars. Data were recorded seven hours (genetic background B), four hours (background A), or five hours (*cis*-1,4-polyisoprene) after the start of the experiment, at timepoints when larvae first showed significant choice behavior. **(b)** Diet consumption, larval weight gain and larval weight gain per consumed diet of *M. melolontha* larvae feeding for five consecutive days on artificial diet with 3% *cis*-1,4-polyisoprene or solvent control. *P*-values refer to a linear mixed effect model in the left panel, and Wilcoxon signed-rank tests in the middle and right panels. N=24-26.

Second, we assessed *M. melolontha* feeding preference between carrot seedlings coated with ecologically relevant concentrations of purified *cis*-1,4-polyisoprene or a solvent control. Five hours after the start of the experiment, larvae preferred to feed on control rather than on *cis*-1,4-polyisoprene supplemented roots (*P*=0.014, binomial test, Fig. 2a), with 69% of the larvae feeding on the control roots.

Third, we measured larval growth and food consumption over five consecutive days in a non-choice assay in which larvae were offered artificial diet supplemented with either an ecologically relevant concentration of *cis*-1,4-polyisoprene or a solvent control. Surprisingly, larvae consumed on average 13% more *cis*-1,4-polyisoprene supplemented diet compared to the control diet (*P*=0.003, linear mixed effect model, Fig. 2b). Nevertheless, larvae lost three-fold more weight within the five days on the *cis*-1,4-polyisoprene supplemented diet (*P*=0.01, Wilcoxon test, Fig. 2b). Consequently, *cis*-1,4-polyisoprene supplementation reduced larval weight gain per mg consumed diet (*P*=0.048, Wilcoxon test, Fig. 2b).

Finally, we specifically tested the effect of pentacyclic triterpenes - which are elevated three-fold in the rubber-depleted *Tk*CPTL1-RNAi compared to NIL lines (Niephaus *et al*., 2019) - on plant performance under herbivory and larval growth using two independent triterpene-reduced *TkOSC*-RNAi lines and their respective triterpene-containing NIL lines (van Deenen *et al*., 2019). Plant resistance and larval weight gain between triterpene-containing and deficient plants did not differ (Supporting Information Figs. S2-S3). To complement the transgenic approach, we assessed whether lupeol, a major *T. koksaghyz* triterpene (Pütter *et al*., 2019), alters *M. melolontha* food consumption and larval growth. Lupeol did not alter diet consumption or larval weight gain when supplemented in ecologically relevant concentrations to artificial diet (Supporting Information Fig. S4).

Taken together, these data provide complementary evidence that *cis*-1,4-polyisoprene deters *M. melolontha* feeding and thereby benefits *T. koksaghyz* under *M. melolontha* attack.

### Natural rubber biosynthesis alters the structure of the *T. koksaghyz* root and rhizosphere microbiome in a herbivore- and wounding-dependent manner

Apart from reducing herbivory, natural rubber is hypothesized to alter the plant microbiome, particularly by sealing wounds and thereby preventing the entry of pathogens. To test this hypothesis, we assessed the root and rhizosphere microbiome of rubber-depleted *Tk*CPTL1-RNAi and the normal rubber content control NIL lines in the A genetic background using plants grown in natural field soil under control, *M. melolontha* herbivory and wounding treatments. We first tested the hypothesis that natural rubber and the damage treatments (wounding or herbivory) alter the microbial diversity. Based on both Shannon and Simpson indices, neither the RNAi silencing nor the damage treatments altered the rhizosphere or root microbial diversity (Supporting Information Fig. S6, Table S2-S5).

Next, we tested whether RNAi silencing and the damage treatments alter the root and rhizosphere microbial structure. In the rhizosphere, the structure of the bacterial but not fungal community was affected by both the plant genotype and the damage treatment (*P*(NIL control – RNAi control)=0.0005 for bacteria and 0.466 for fungi; P(NIL control – NIL herbivory or NIL wounding) = 0.0005 for bacteria and >0.5 for fungi; Table 1, Supporting Information Figs. S7-S8). Interestingly, the effect of the damage treatment depended on the plant genotype: In NIL plants, both herbivory and wounding influenced the structure of the rhizosphere bacterial community, whereas in *Tk*CPTL1-RNAi plants, only herbivory had such an effect (pMCMC=0.01). In contrast, in NIL plants, neither herbivory nor wounding affected the structure of the rhizosphere fungal community (pMCMC>0.47), whereas in *Tk*CPTL1-RNAi plants, both herbivory and wounding altered the rhizosphere fungal community (pMCMC<0.03, Table 1, Supporting Information Figs. S9-S10). These changes were also reflected in the microbial gene pool of the rhizosphere obtained from shotgun metagenomic sequencing, which was altered by both the plant genotype (Table 1, Fig. 3a) and the treatment within each plant genotype (Table 1, Supporting Information Fig. S11).

**Table 1.**
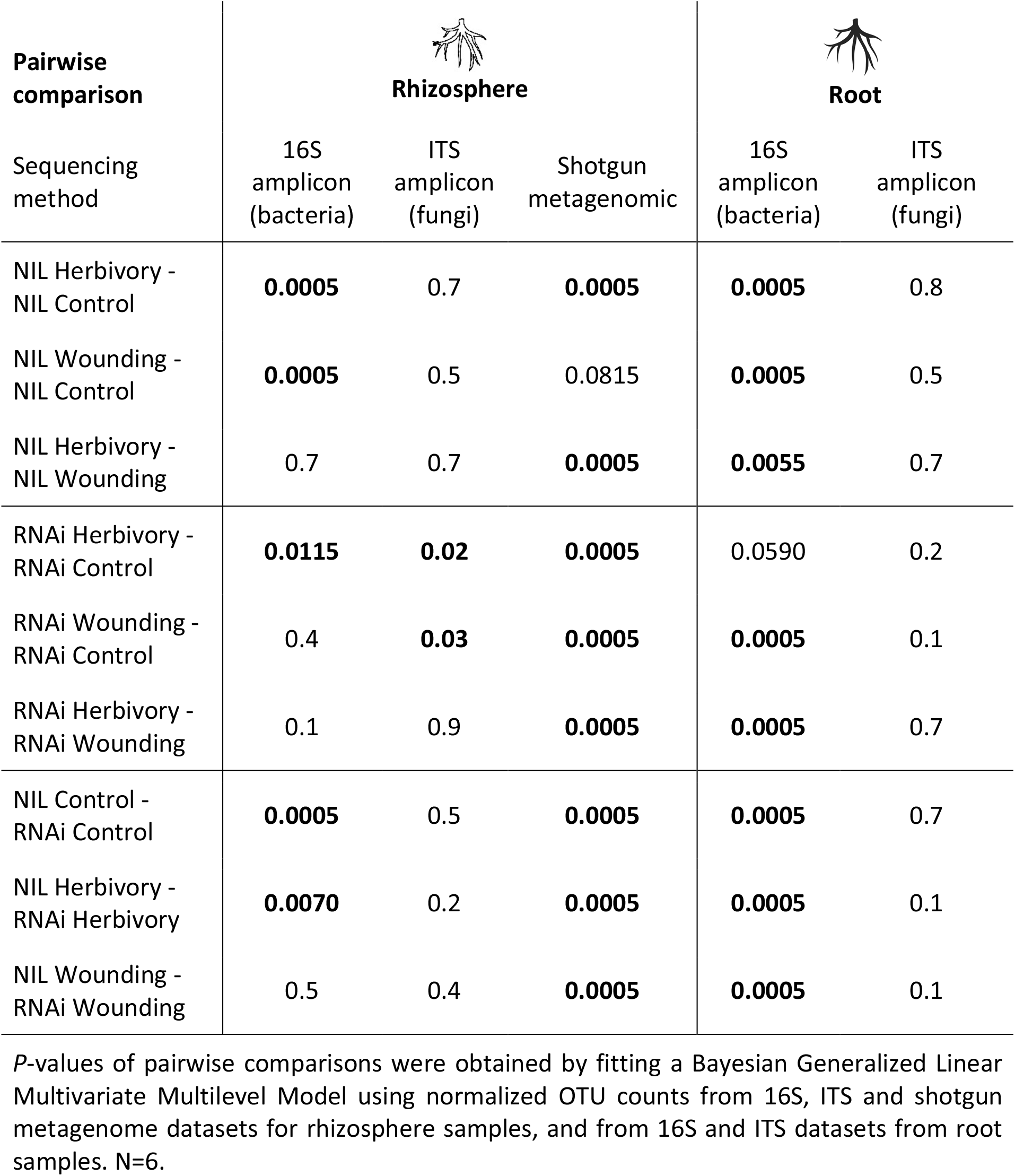
*P*-values (pMCMC) of multivariate analysis investigating the structure of the rhizosphere and root microbiome of rubber-deficient *Tk*CPTL1-RNAi and normal rubber content NIL *Taraxacum koksaghyz* plants growing in natural field soil under control conditions, *Melolontha melolontha* herbivory or mechanical wounding.

| Pairwise comparison | Rhizosphere |  |  | Root |  |
| --- | --- | --- | --- | --- | --- |
|  | 16S amplicon (bacteria) | ITS amplicon (fungi) | Shotgun metagenomic | 16S amplicon (bacteria) | ITS amplicon (fungi) |
| NIL Herbivory - NIL Control | **0.0005** | 0.7 | **0.0005** | **0.0005** | 0.8 |
| NIL Wounding - NIL Control | **0.0005** | 0.5 | 0.0815 | **0.0005** | 0.5 |
| NIL Herbivory - NIL Wounding | 0.7 | 0.7 | **0.0005** | **0.0055** | 0.7 |
| RNAi Herbivory - RNAi Control | **0.0115** | **0.02** | **0.0005** | 0.0590 | 0.2 |
| RNAi Wounding - RNAi Control | 0.4 | **0.03** | **0.0005** | **0.0005** | 0.1 |
| RNAi Herbivory - RNAi Wounding | 0.1 | 0.9 | **0.0005** | **0.0005** | 0.7 |
| NIL Control - RNAi Control | **0.0005** | 0.5 | **0.0005** | **0.0005** | 0.7 |
| NIL Herbivory - RNAi Herbivory | **0.0070** | 0.2 | **0.0005** | **0.0005** | 0.1 |
| NIL Wounding - RNAi Wounding | 0.5 | 0.4 | **0.0005** | **0.0005** | 0.1 |

*P*-values of pairwise comparisons were obtained by fitting a Bayesian Generalized Linear Multivariate Multilevel Model using normalized OTU counts from 16S, ITS and shotgun metagenome datasets for rhizosphere samples, and from 16S and ITS datasets from root samples. N=6.

**Figure 3.**
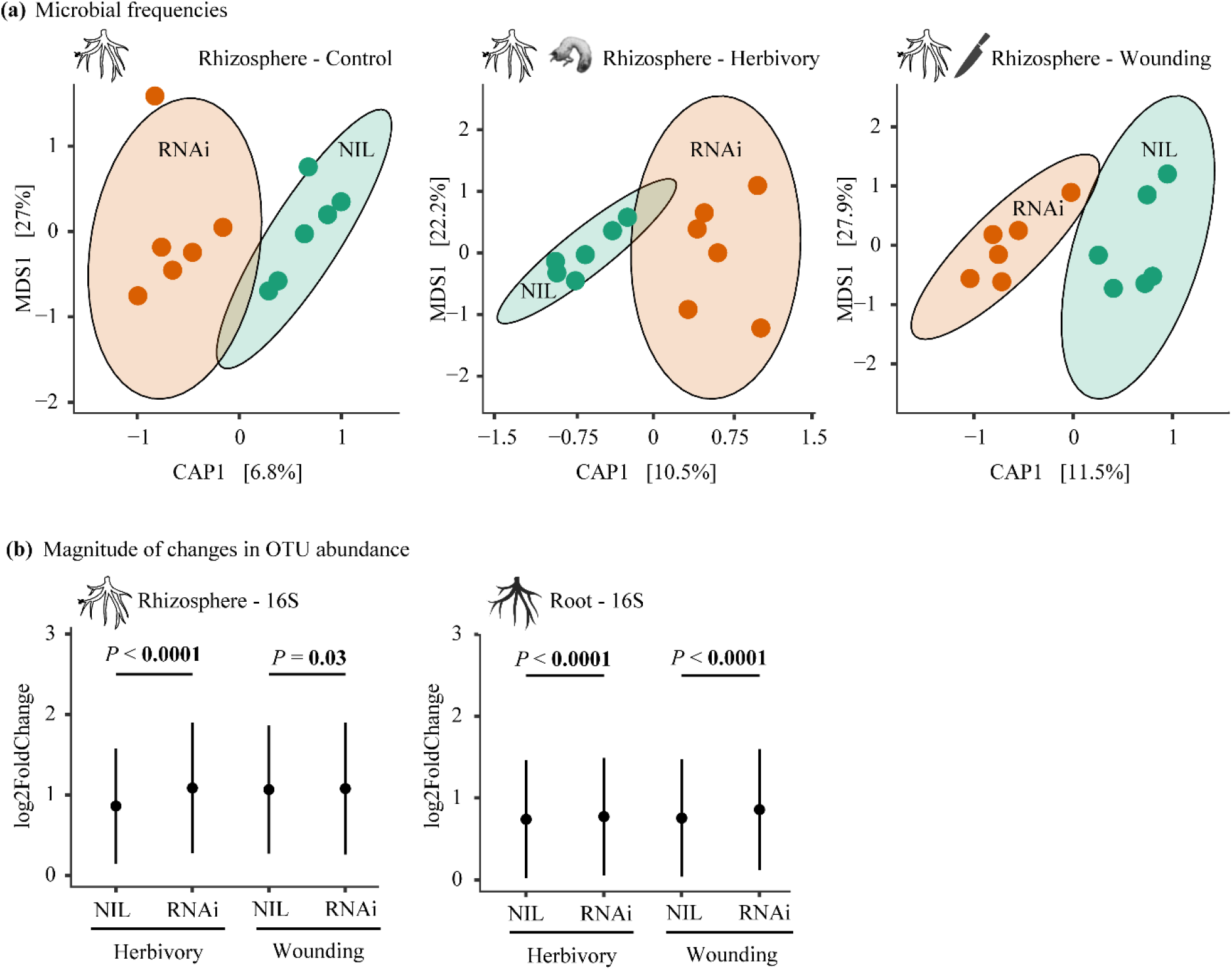
Abolishing the biosynthesis of *cis*-1,4-polyisoprene through RNAi alters the *Taraxacum koksaghyz* microbiome in an herbivore- and wounding-dependent manner. **(a)** Canonical analysis of principal (CAP) coordinates ordination (Bray–Curtis distance matrix) on microbial gene frequencies from rhizosphere samples obtained from shotgun metagenomic sequencing, reporting the effect of plant genotype (NIL, *Tk*CPTL1-RNAi, different colors) under control, herbivory and mechanical wounding. Percentages in parentheses report the variance explained by the respective axis. Statistical analysis on the pairwise comparisons are found in Table 1 under “shotgun metagenomic”). **(b)** The magnitude of changes in the abundance of each bacterial OTU (absolute log2 fold changes) upon wounding and herbivory differed between *Tk*CPTL1-RNAi and NIL plants. Data was analyzed using a linear mixed-effects model for bacterial rhizosphere samples (χ2=1900, df=3, *P*<0.001) and bacterial root samples (χ2=250.64, df=3, *P*<0.001). *P*-values of pairwise comparisons were adjusted using the FDR method. For fungal OTUs see also Supporting Information Fig. S12 and Table S5. N=6.

Similar to the rhizosphere, in the plant roots, the bacterial but not the fungal community structure was altered by the plant genotype, treatment and their interaction (Supporting Information Figs. S7-S10). Specifically, in NIL plants, both herbivory and wounding influenced the structure of the root bacterial community (pMCMC<0.001), while in *Tk*CPTL1-RNAi plants only wounding had this effect (pMCMC<0.001) (Table 1, Supporting Information Fig. S9). None of the treatments altered the structure of the fungal community in either plant genotype (Table 1, Supporting Information Fig. S10).

Since multivariate analysis only reveals differences in microbial community structure, but not in the extent of these changes, we calculated the magnitude of change in the rhizosphere and root microbiomes (absolute log2 Fold Changes) upon wounding or herbivory in each plant genotype. In the rhizosphere, the bacterial microbiota showed a larger magnitude of change upon both herbivory and wounding in *Tk*CPTL1-RNAi than NIL plants (Fig. 3b, Supporting Information Table S5), whereas the fungal microbiota exhibited a higher magnitude of change upon herbivory but lower magnitude of change upon wounding in *Tk*CPTL1-RNAi compared to NIL plants (Supporting Information Fig. S12, Table S5). In the roots, a similar pattern was observed: the bacterial community exhibited a higher magnitude of change upon herbivory or wounding in *Tk*CPTL1-RNAi than NIL plants (Fig. 3b, Supporting Information Table S6), while the fungal community was unaffected by the plant genotype (Supporting Information Fig. S12, Table S5).

### Abolishing biosynthesis of natural rubber did not increase the microbial colonization or pathogen load in *T. koksaghyz* roots upon injury

As both the abundance of *cis*-1,4-polyisoprene (plant genotype) and herbivory or mechanical wounding treatment shaped the structure of the rhizosphere and root microbiota, we used four approaches to test the hypothesis that biosynthesis of the dominant metabolite of natural rubber restricts the entry of potentially pathogenic microorganisms into the roots.

First, we used shotgun metagenomics to test whether plant genotype and treatment influenced the microbial load in plant roots (i.e. the percentage of sequence reads that did not align to the plant genome relative to the reads that aligned to the host plant). While the percentage of microbial reads in the roots showed a tendency to be altered by the treatment (*P*=0.051, linear model), these changes were independent of plant genotype (*P*>0.5, linear model, Fig. 4a).

**Figure 4.**
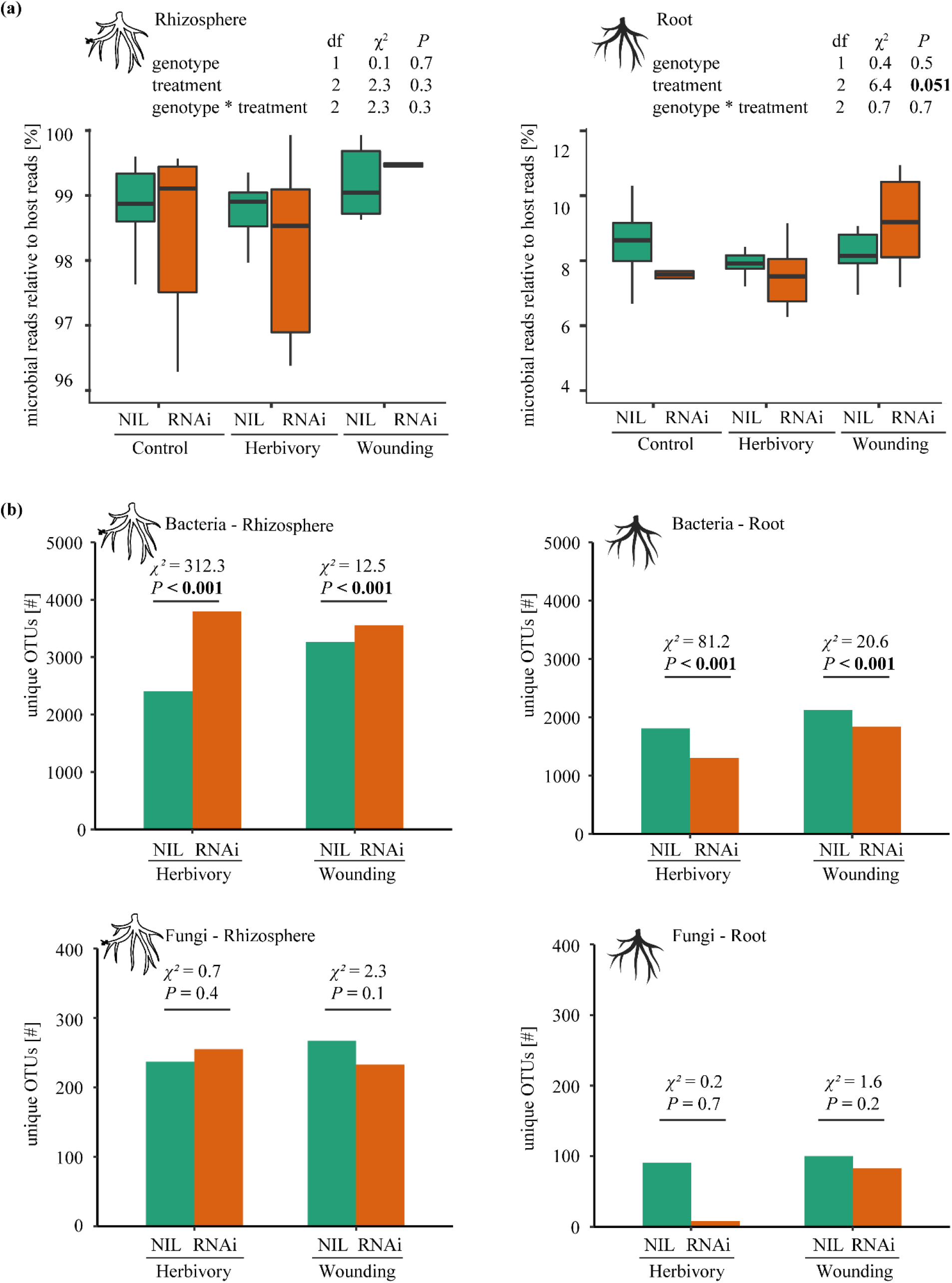
**(a)** Abolishing the biosynthesis of *cis*-1,4-polyisoprene through RNAi does not increase microbial colonization or pathogen load in *Taraxacum koksaghyz* roots. **(a)** The percentage of microbial reads in rhizosphere and root samples for each treatment (control, herbivory or wounding) and plant genotype (rubber-deficient *Tk*CPTL1-RNAi, “RNAi”, and normal rubber content NIL, “NIL”, plants). *P*-values of linear models are shown on top of the panels. N=6. **(b)** The number of bacterial and fungal OTUs in rhizosphere and roots of rubber-deficient *Tk*CPTL1-RNAi and normal rubber content NIL *T. koksaghyz* plants that are unique to the herbivore and wounding treatment (i.e., not found under control conditions). Statistics refer to χ^2^ tests. N=6.

Second, if natural rubber restricts the colonization of roots, we would expect a higher number of OTUs unique to the herbivore or wounding treatment (i. e. not found under control conditions) in rubber-depleted *Tk*CPTL1-RNAi compared to normal rubber content NIL plants, particularly in roots. Contrary to our expectations, the numbers of bacterial OTUs unique to the herbivore and wounding treatments were lower in the roots but higher in the rhizosphere in the *Tk*CPTL1-RNAi compared to NIL plants (Fig. 4b). The number of fungal OTUs unique to the herbivore or wounding treatment did not differ between *Tk*CPTL1-RNAi and NIL plants (Fig. 4b).

Third, we assessed whether taxa that are differentially abundant between NIL and *Tk*CPTL1-RNAi plants under wounding or herbivory are pathogens. Although we found taxa that were differentially abundant in each *plant_genotype*treatment* combination, none of those taxa appeared to be agents of plant diseases (Supporting Information Notes S1). Also, the pathogens and endophytes (pathogens: *Botrytis spp;* endophytes: *Curtobacterium sp*., *Leifsonia sp*., *Methylobacterium sp*., *Microbacterium spp*.) that had been recorded in the field for these *T. koksaghyz* lines (personal communication Fred Eickmeyer) and were present in our dataset, were similarly abundant across the damage treatments and plant genotypes in both root and rhizosphere samples (Supporting Information Notes S1, Tables S6-S12).

Fourth, to obtain more insights into the function of the plant genotype-dependent taxonomic changes upon wounding and herbivory, we tested whether the changes in the microbial gene frequencies of each *plant_genotype*treatment* combination obtained by shotgun metagenomic sequencing compared to the control treatment of the respective plant genotype were significantly different from random. We found a higher number of differentially abundant genes in all the groups compared to what we would expect by chance (Supporting Information Table S14). Within each *plant_genotype*treatment* combination, we classified each differentially abundant gene according to its likely functional role within the community. Most of the differentially abundant genes were related to metabolic functions (Supporting Information Table S15). However, we also found a group of differentially abundant genes that have been previously reported in plant pathogens as pathogenicity or virulence factors. The number of these pathogen-related genes that were enriched in the wounding or herbivory treatment was at least three times higher in NIL than in *Tk*CPTL1-RNAi plants (Table 2).

**Table 2.** Microbiome functional analysis.

| Contrast | Microbial genes |
| --- | --- |
| **NIL Herbivory -**<br>**NIL Control** | Acetyl-coenzyme A synthetase<sup>1</sup><br>ATP-dependent RNA helicase DeaD<sup>2</sup><br>ATP-dependent RNA helicase SrmB<sup>3</sup><br>Beta-N-acetylglucosaminidase/beta-glucosidase<sup>4</sup><br>Beta-xylosidase<sup>5</sup><br>Levanase<sup>6</sup><br>Mannitol 2-dehydrogenase<sup>7</sup><br>Multidrug efflux pump subunit AcrB<sup>8</sup><br>Sensor protein KdpD<sup>9</sup><br>Translation initiation factor IF-2<sup>10</sup> |
| **NIL Wounding -**<br>**NIL Control** | Acetyl-coenzyme A synthetase<sup>1</sup><br>Acyl-CoA dehydrogenase<sup>11</sup><br>ATP-dependent RNA helicase DeaD<sup>2</sup><br>ATP-dependent RNA helicase RhlE<sup>12</sup><br>Beta-xylosidase<sup>5</sup><br>Bifunctional chorismate mutase/prephenate dehydratase<sup>13</sup><br>Cystathionine gamma-synthase<sup>14</sup><br>Extracellular serine protease<sup>15</sup><br>Glucans biosynthesis protein G<sup>16</sup><br>Nitrilotriacetate monooxygenase component A<sup>17</sup> |
| **RNAi Herbivory -**<br>**RNAi Control** | Acyl-CoA dehydrogenase<sup>11</sup><br>ATP-dependent RNA helicase SrmB<sup>3</sup><br>putative sensor histidine kinase TcrY<sup>18</sup> |
| **RNAi Wounding -**<br>**RNAi Control** | - |

Microbial genes previously reported in the literature to be associated with pathogenicity or virulence that were found significantly more abundant in the rhizosphere of treated plants (herbivory or wounding) than the respective control group. "NIL" indicates plants with normal rubber content and "RNAi" indicates rubber-depleted plants. <sup>1</sup>(Gu *et al.*, 2019), <sup>2</sup>(Li *et al.*, 2008), <sup>3</sup>(Granato *et al.*, 2016), <sup>4</sup>(Huang *et al.*, 2021), <sup>5</sup>(Guzha *et al.*, 2022), (Versluys *et al.*, 2018), <sup>7</sup>(Jennings *et al.*, 1998), <sup>8</sup>(Burse *et al.*, 2007), <sup>9</sup>(Yang *et al.*, 2018), <sup>10</sup>(Zhang *et al.*, 2013), <sup>11</sup>(Ruswandi *et al.*, 2005), <sup>12</sup>(Hausmann *et al.*, 2021), <sup>13</sup>(Kim *et al.*, 2020), <sup>14</sup>(Fu *et al.*, 2013), <sup>15</sup>(Figaj *et al.*, 2019), <sup>16</sup>(Page *et al.*, 2001), <sup>17</sup>(Agarwal *et al.*, 2021), <sup>18</sup>(Cai *et al.*, 2017)

Taken together, these data provide evidence that biosynthesis of natural rubber alters the microbiome in a herbivore- and wounding-dependent manner, but that these changes are not associated with an increased pathogen load in the rubber-depleted plants.

## Discussion

Laticifers – which are among the most common secretory structures in flowering plants - have long been hypothesized to mediate both plant-herbivore and plant-microbe interactions. Yet, experimental evidence for either of these functions is scarce. Here, we provide evidence that the *cis*-1,4-polyisoprene, the major constituent of natural rubber, helps to protect *T. koksaghyz* against the generalist root feeder *M. melolontha*. Furthermore, the biosynthesis of *cis*-1,4-polyisoprene alters the microbial colonization of *T. koksaghyz* roots and rhizosphere in a herbivore- and wounding-dependent manner. Thereby, our study sheds light on the long-standing mystery of why some plants produce natural rubber.

### *Cis*-1,4-polyisoprene reduces herbivory

Natural rubber has long been hypothesized to be defensive against herbivores, but experimental evidence for this notion is scarce. Combining genetic and chemical supplementation approaches *in* and *ex planta*, we obtained parallel lines of evidence that natural rubber reduces herbivory. First, rubber-deficient *Tk*CPTL1-RNAi lines silenced in *cis*-1,4-polyisoprene biosynthesis were preferred by *M. melolontha* and lost more biomass upon herbivory than normal rubber content NIL lines. While the benefits of natural rubber biosynthesis on plant performance under herbivory were weaker in genetic background A than B, likely because *M. melolontha* herbivory was less severe in the former, the preference of *M. melolontha* for rubber-deficient plants was consistent in both transgenic lines. Second, purified *cis*-polyisoprene reduced both larval weight gain and the attractiveness of the food. The observed temporal patterns in the choice of *M. melolontha* – with first a gradual increase of the number of larvae on the polyisoprene-deficient side, followed by re-distribution of the larvae in an undirected manner– is typical for *M. melolontha* responding to deterrent compounds (Huber *et al*., 2021); this temporal pattern likely emerges because feeding reduces the attractiveness of the food. Third, *T. koksaghyz* triterpenes - which are elevated in rubber-depleted *Tk*CPTL1-RNAi lines (Niephaus *et al*., 2019) - did not alter herbivore growth and plant resistance.

Our results are in line with a recent study showing that the addition of a *trans*-1,4-polyisoprene to artificial diet deters feeding and larval growth of the longhorn beetle that naturally feeds on the *trans*-1,4-polyisoprene-producing *Eucommia* trees (Pan *et al*., 2015). As specialized metabolites may, however, function differently in isolation than when surrounded by their native matrix (Niemeyer, 2009; Chen *et al*., 2020; Huber *et al*., 2021), *in planta* experiments as carried out in our study are critical to infer ecological functions (Steppuhn *et al*., 2004; Huber *et al*., 2016b; Erb, 2018). Future work that elucidates how the branching structure, molecular weight and concentration of *cis*-1,4-polyisoprene affects plant-environment interactions would provide exciting insights into why this polymer is so variable in nature.

Several mechanisms may contribute to the defensive properties of natural rubber under herbivore attack. Natural rubber is thought to contribute to the stickiness of latex, and thereby may trap insects or immobilize their mouth parts (Agrawal & Konno, 2009). *M. melolontha* is too large to be entrapped by latex, however, and we did not observe that the larva’s mouth parts were glued together. Nevertheless, it is possible that smaller insects could be affected by the stickiness of natural rubber. Alternatively, natural rubber may act as a matrix that facilitates the distribution and penetrance of other latex specialized metabolites inside the herbivore. Indeed, *cis*-polyisoprenes help to disperse antimicrobial triterpenes *in vitro* (Salomé-Abarca *et al*., 2021). In our experiment, addition of *cis*-polyisoprene to artificial diet was sufficient to reduce larval weight gain in the absence of other defensive metabolites, indicating that the polymer at least partially acts by itself. Furthermore, addition of *cis*-1,4-polyisoprene reduced *M. melolontha* weight gain per unit of consumed diet. We thus hypothesize that *cis*-1,4-polysioprene, being chemically inert, impedes nutrient uptake possibly by binding to nutrients or by agglutinating the epithelium of the herbivore’s midgut. Experiments that assess herbivore physiology and localize *cis*-1,4-polyisoprene inside the organism are needed to elucidate the mode of action of natural rubber in herbivore defense.

### *Cis*-1,4-polyisoprene alters the microbiome but not the pathogen load below ground

Apart from herbivore defense, laticifers and in particular natural rubber are hypothesized to mediate plant-microbe interactions (Agrawal & Konno, 2009). We found that abolishing the biosynthesis of *cis*-1,4-polysioprene altered the structure of both the root and the rhizosphere microbiome in non-infested, undamaged plants. This is similar to recent work showing that silencing the biosynthesis of other specialized metabolites, for instance benzoxazinoids, alters the root and rhizosphere microbiome (Stringlis *et al*., 2018; Cotton *et al*., 2019; Koprivova *et al*., 2019; Jacoby *et al*., 2021; Pang *et al*., 2021). At least two mechanisms may account for the observed rubber-associated changes in the microbiome structure in the absence of plant damage:

First, natural rubber may structure the microbiota through direct exposure, as on the one hand endophytes may inhabit laticifers (Gunawardana *et al*., 2015) and on the other hand natural rubber is released to the soil when roots decay. We, however, did not observe any changes in the abundance of known *T. koksaghyz* endophytes with variation in plant rubber content. Second, abolishing the biosynthesis of *cis*-1,4-polysioprene through RNAi may alter the microbial structure through changes in metabolic fluxes, particularly in triterpene biosynthesis (Niephaus *et al*., 2019). Triterpenes may promote or inhibit microbial growth (Pacheco *et al*., 2012), and disrupting triterpene biosynthesis in *A. thaliana* altered the root microbiome (Huang *et al*., 2019). Assessing the effect of natural rubber on the growth of selected microbes, and complementing the rhizosphere of rubber-deficient plants with relevant *cis*-1,4-polyisoprene concentrations could help to differentiate among these possibilities.

The major role of natural rubber in plant-microbe interactions is hypothesized to be wound sealing, thereby preventing the entry of pathogens. We obtained mixed evidence for this hypothesis: on the one hand, abolishing the biosynthesis of *cis*-1,4-polysioprene altered the structure of the rhizosphere and root microbiota in a herbivore- and wounding-dependent manner. Furthermore, rubber-deficient *Tk*CPTL1-RNAi plants exhibited a larger magnitude of change in the root microbiota upon wounding or herbivory compared to normal rubber content NIL plants. These results support the role of natural rubber in wound sealing, and emphasize that plant-microbe and plant-herbivore interactions should be studied in concert when assessing the function of specialized metabolites (Hu *et al*., 2018; Kudjordjie *et al*., 2021).

On the other hand, silencing the biosynthesis of *cis*-1,4-polysioprene did not alter the percentage of microbial reads in plant roots upon damage, nor increase the number of OTUs in roots unique to damaged plants, nor increase the number of pathogenicity-related microbial genes that are enriched in roots upon damage. These finding contrast with recent *in vitro* assays showing that *cis*-1,4-polyisoprene forms a physical barrier to bacteria and to a lower extent also to fungi including *Botrytis* spp. (Salomé-Abarca *et al*., 2021). The different conditions *in vitro* and *in planta*, as well as variation in the amount, molecular weight and branching structure of *cis*-1,4-polyisoprene may contribute to differences in the inferred role of this polymer in pathogen defense. Future experiments that assess the performance of rubber-depleted and normal rubber content plants in combination with manipulation of the soil microbiota similar to (Hu *et al*., 2018; Kudjordjie *et al*., 2021) will be critical to improve our understanding of the role of natural rubber in pathogen defense.

Taken together, we have shown that biosynthesis of *cis*-1,4-polysioprene reduces herbivory and structures the microbiome in an herbivore- or wounding-dependent manner. Therefore, latex mediates both plant-herbivore and plant-microbe interactions, highlighting the role of plant specialized metabolites and secretory structures in shaping the complex trophic interactions in nature.

## Data availability

The data that support the findings of this study are openly available in NCBI SRA under the BioProject numbers PRJNA779274 (16S amplicon metagenomics), PRJNA779290 (ITS amplicon metagenomics) and PRJNA779369 (shotgun metagenomics). Public code repository can be found here: https://github.com/amalacrino/dandelion_microbiome.

## Acknowledgements

We thank Eva Niephaus and Kristina Unland for providing transgenic seed material and Zoe Bont, Christelle Robert, Matthias Erb and Markus Dönz for providing the first batch of *M. melolontha* larvae. We thank Otto Fäth, Frederike Sasse and Mark Winzen for their help during the collecting of the second batch of larvae. We are grateful to Fred Eickmeyer (ESKUSA GmbH, Parkstetten, Germany) for providing field soil and information on pathogens in the field. We thank Sascha Ahrens, Pascal Poweleit, Vincent Benninghaus, Joachim Rikus and Daniela Ahlert for experimental support, and Martin Schäfer for fruitful discussions.

## Author contribution

MH conceived the study. MH, LB, NvD and BM designed the experiments. LB performed experiments. LB, AM and MH analyzed data. MH, CSG and DP supervised students. DP, JG, MH and SX contributed resources. LB, AM and MH wrote the initial draft, and all authors contributed to its final version.

## The following Supporting Information is available for this article

**Fig. S1** Choice experiments *ex planta*, including all timepoints.

**Fig. S2** Biomass accumulation of triterpene-reduced plants under herbivory (Line 2)

**Fig. S3** Biomass accumulation of triterpene-reduced plants under herbivory (Line 3)

**Fig. S4** Larval weight gain and diet consumption of larvae feeding on lupeol supplemented diet

**Fig. S5** Resistance of rubber-deficient RNAi and NIL plants (A line) grown in natural field soil.

**Fig. S6** Diversity analysis (Shannon and Simpson indexes) for 16S ITS rhizosphere and root

**Fig. S7** CAP on bacterial metabarcoding reporting effect of genotype on rhizosphere and root

**Fig. S8** CAP on fungal metabarcoding reporting effect of genotype on rhizosphere and root

**Fig. S9** DCA on bacterial metabarcoding reporting effect of treatment

**Fig. S10** DCA on fungal metabarcoding reporting effect of treatment

**Fig. S11** CAP on microbial gene content on rhizosphere reporting the effect of treatment

**Fig. S12** Magnitude of changes in abundance for fungal OTU

**Table S1** Diversity analysis. Model coefficients Shannon diversity

**Table S2** Diversity analysis. Model coefficients Simpson diversity

**Table S3** Diversity analysis. Pairwise comparisons Shannon index

**Table S4** Diversity analysis. Pairwise comparisons Simpson index

**Table S5** Magnitude of changes in abundance of bacterial and fungal OTU

**Table S6** Number of observed vs random OTUs in rhizosphere

**Table S7** Number of observed OTUs that are differently abundant in rhizosphere 16S data

**Table S8** Number of observed OTUs that are differently abundant in rhizosphere ITS data

**Table S9** Number of observed vs random OTUs in roots

**Table S10** Number of observed OTUs that are differently abundant in roots 16S data

**Table S11** Relative abundance of bacterial genus in plant rhizosphere

**Table S12** Relative abundance of bacterial genus in plant roots

**Table S13** Functional analysis reporting the number of observed genes differently abundant

**Table S14** List of genes and corresponding group stating a higher abundance

**METHODS S1** Plant material and growth conditions

**METHODS S2** Identification of transgenic rubber-depleted plants

**METHODS S3** Isolation of pure *cis*-1,4-polyisoprene

**METHODS S4** Details on triterpene experiments

**METHODS S5** Origin and storage of soil used for microbiome experiment

**METHODS S6** Details on sample handling of microbiome study and statistical analysis

**NOTES S1** Taxa analysis

**REFERENCES**

