## Supporting Information for "Natural rubber reduces herbivory and alters the microbiome below ground"

The following Supporting Information is available for this article:

**Fig. S1** Choice experiments *ex planta*, including all timepoints.

**Fig. S2** Biomass accumulation of triterpene-reduced plants under herbivory (Line 2)

**Fig. S3** Biomass accumulation of triterpene-reduced plants under herbivory (Line 3)

**Fig. S4** Larval weight gain and diet consumption of larvae feeding on lupeol supplemented diet

**Fig. S5** Resistance of rubber-deficient RNAi and NIL plants (A line) grown in natural field soil

**Fig. S6** Diversity analysis (Shannon and Simpson indexes) for 16S ITS rhizosphere and root

**Fig. S7** CAP on bacterial metabarcoding reporting effect of genotype on rhizosphere and root

**Fig. S8** CAP on fungal metabarcoding reporting effect of genotype on rhizosphere and root

**Fig. S9** DCA on bacterial metabarcoding reporting effect of treatment

**Fig. S10** DCA on fungal metabarcoding reporting effect of treatment

**Fig. S11** CAP on microbial gene content on rhizosphere reporting the effect of treatment

**Fig. S12** Magnitude of changes in abundance for fungal OTU

**Table S1** Diversity analysis. Model coefficients Shannon diversity

**Table S2** Diversity analysis. Model coefficients Simpson diversity

**Table S3** Diversity analysis. Pairwise comparisons Shannon index

**Table S4** Diversity analysis. Pairwise comparisons Simpson index

**Table S5** Magnitude of changes in abundance of bacterial and fungal OTU

**Table S6** Number of observed vs random OTUs in rhizosphere

**Table S7** Number of observed OTUs that are differently abundant in rhizosphere 16S data

**Table S8** Number of observed OTUs that are differently abundant in rhizosphere ITS data

**Table S9** Number of observed vs random OTUs in roots

**Table S10** Number of observed OTUs that are differently abundant in roots 16S data

**Table S11** Relative abundance of bacterial genus in plant rhizosphere

**Table S12** Relative abundance of bacterial genus in plant roots

**Table S13** Functional analysis reporting the number of observed genes differently abundant

**Table S14** List of genes and corresponding group stating a higher abundance

**METHODS S1** Plant material and growth conditions

**METHODS S2** Identification of transgenic rubber-depleted plants

**METHODS S3** Isolation of pure *cis*-1,4-polyisoprene

**METHODS S4** Details on triterpene experiments

**METHODS S5** Origin and storage of soil used for microbiome experiment

**METHODS S6** Details on sample handling of microbiome study and statistical analysis

**NOTES S1** Taxa analysis

**REFERENCES**

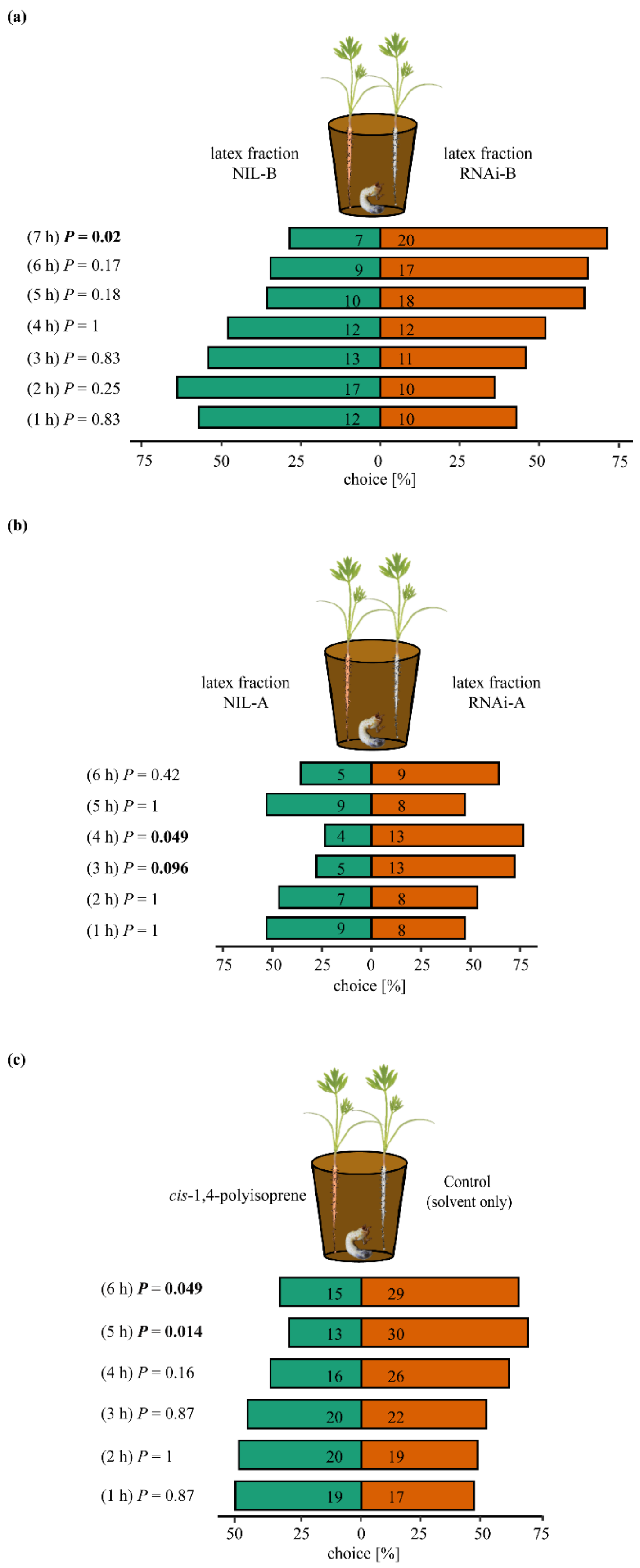


**Fig. S1** Choice of *M. melolontha* between *cis*-1,4-polyisoprene-containing and -lacking food across time. Carrot seedlings were supplemented with (**a**) the whole latex fraction of *T. koksaghyz* rubber-depleted *Tk*CPTL1-RNAi versus normal rubber content NIL plants (B-line; N=28 pairs) (**b**) the whole latex fraction of *T. koksaghyz* rubber-depleted *Tk*CPTL1-RNAi versus normal rubber content NIL plants (A line; N=23 pairs) or **(c)** isolated *cis*-1,4-polyisoprene from *T. koksaghyz* versus solvent only as control. Larvae were allowed to move within the beakers and position was recorded for several time points. N=50 pairs. Numbers in bars indicate absolute number of active larvae per timepoint. *P*-values of binomial tests are shown.

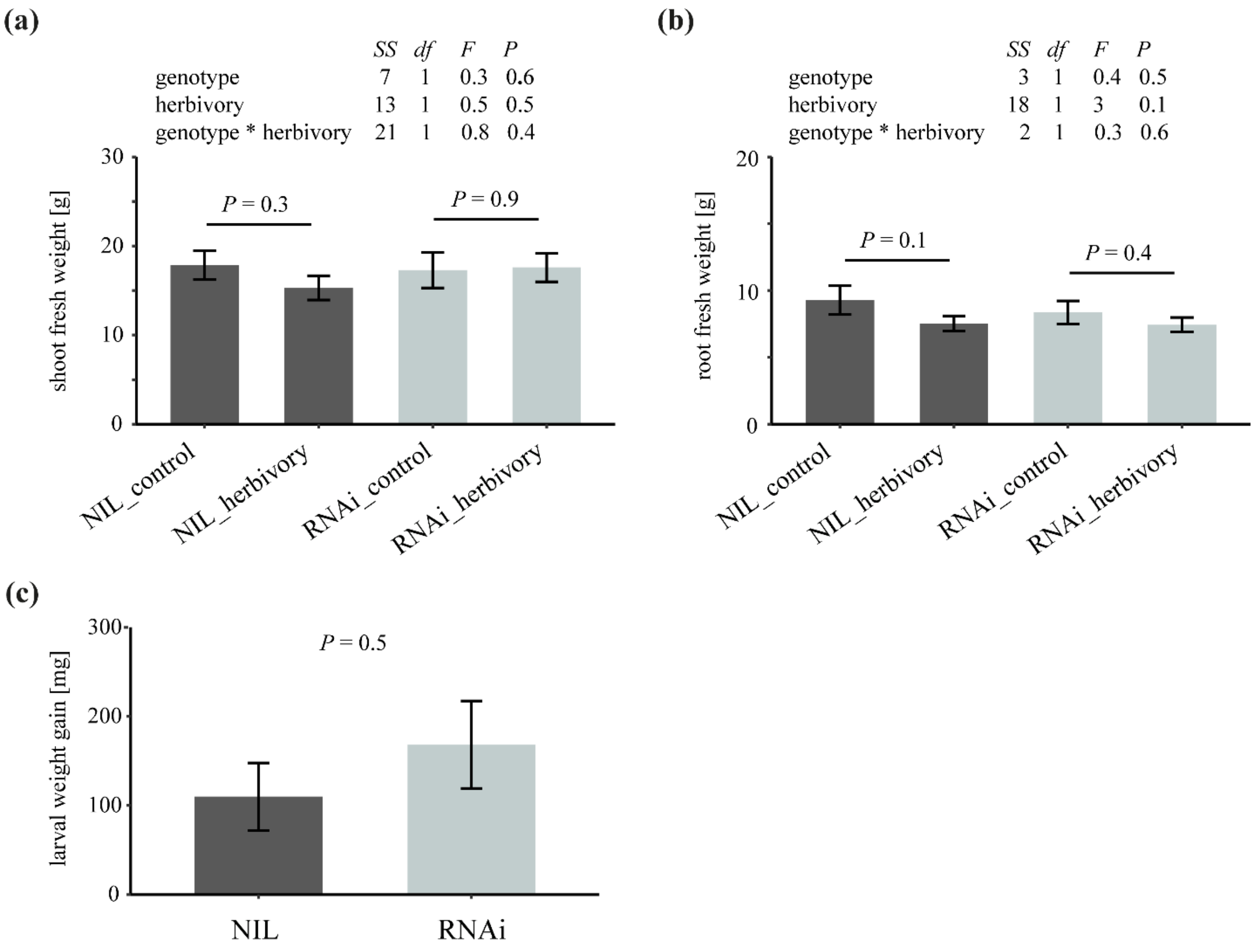


**Fig. S2** Abolishing triterpene biosynthesis did not alter susceptibility of *T. koksaghyz* to *M. melolontha* herbivory in the *TkOSC*-L2 line. **(a)** shoot and **(b)** root fresh mass of triterpene reduced *TkOSC*-RNAi and triterpene-containing NIL plants after nine days of *M. melolontha* herbivory. *P*-values of linear models are displayed. *P*-values of pairwise comparisons refer to FDR corrected values upon extraction with the function emmeans. N=10. **(c)** Larval weight gain after nine days of feeding on NIL or RNAi plants. *P*-values refer to Wilcoxon signed-rank tests. SS=sum squares, df=degrees of freedom, N=10.

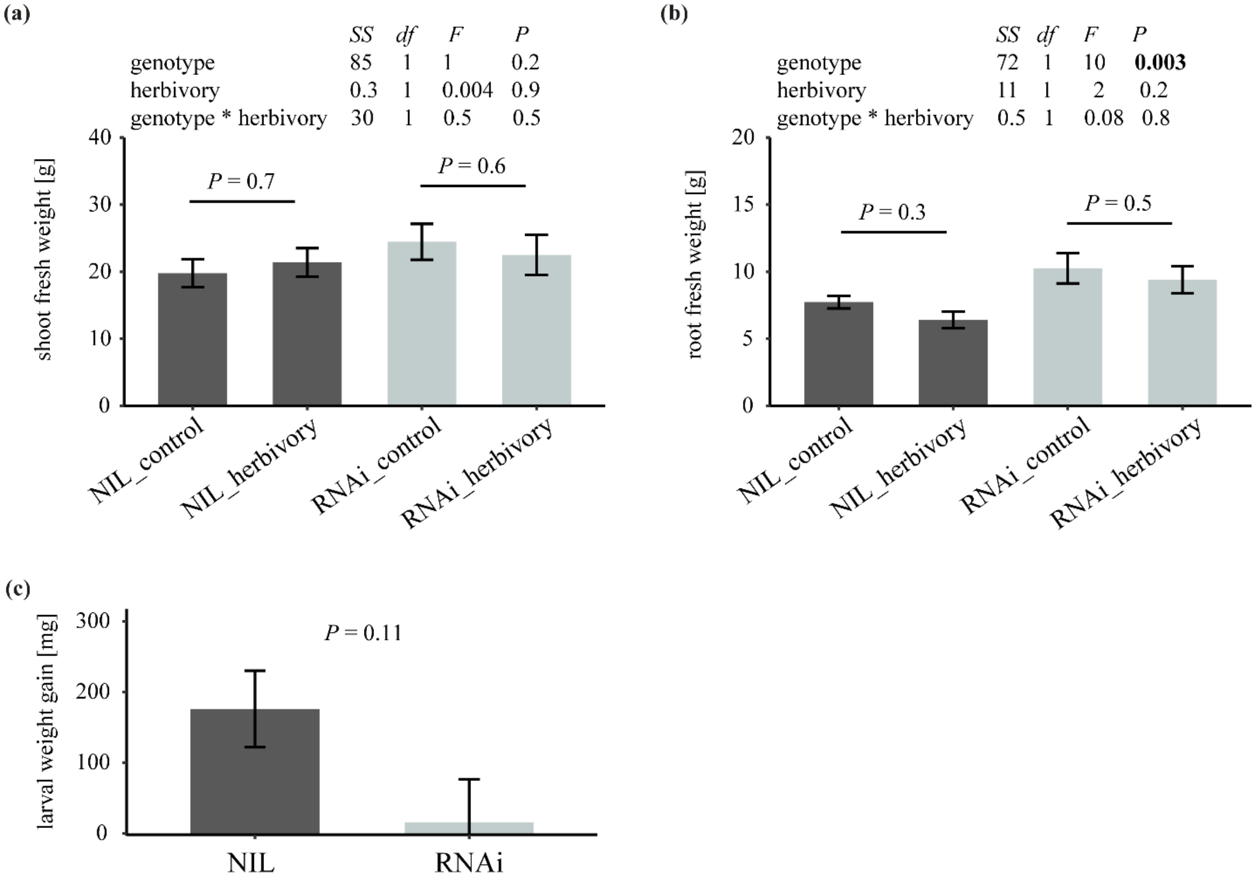


**Fig. S3** Abolishing triterpene biosynthesis did not alter susceptibility of *T. koksaghyz* to *M. melolontha* herbivory in the *TkOSC*-L3 line. **(a)** Shoot and **(b)** root fresh mass of triterpene reduced *TkOSC*-RNAi and triterpene-containing NIL plants upon nine days of *M. melolontha* herbivory. *P*-values of linear models are displayed. *P*-values of pairwise comparisons refer to FDR corrected values upon extraction with the function emmeans. N=9-10. **(c)** Larval weight gain after nine days of feeding on NIL or RNAi plants. *P*-values refer to Wilcoxon signed-rank tests. SS=sum squares, df=degrees of freedom, N=9.

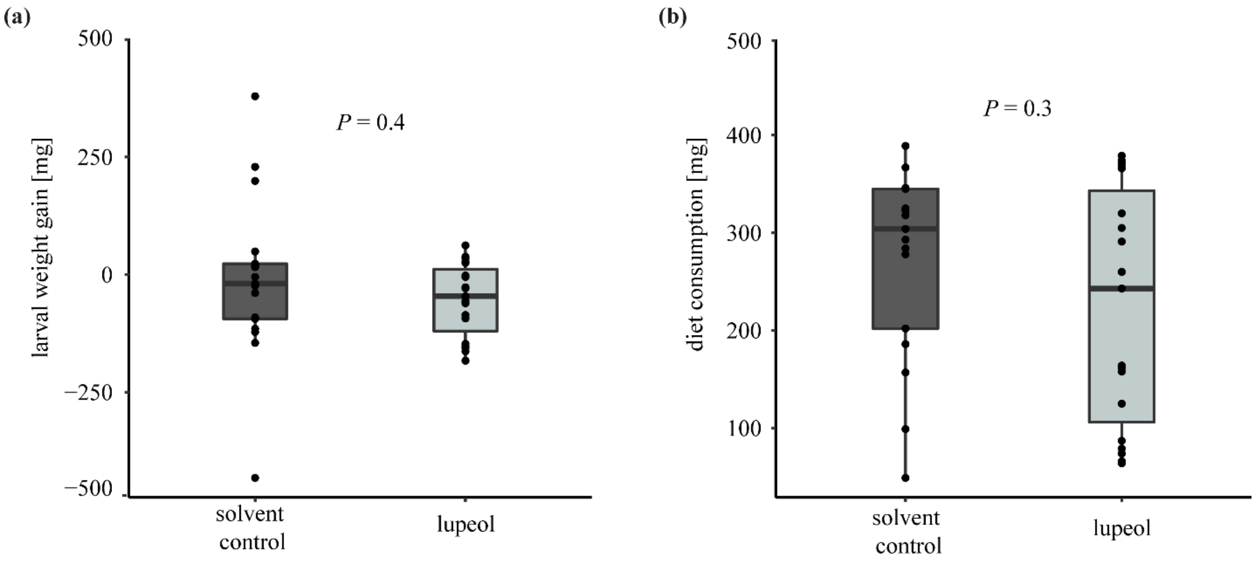


**Fig. S4** Supplementation of lupeol in ecologically relevant concentration to diet (0.1 % based on fresh cube weight) did not affect *M. melolontha* **(a)** weight gain nor **(b)** diet consumption. *P*-values refer to Wilcoxon signed-rank tests. N≥17.

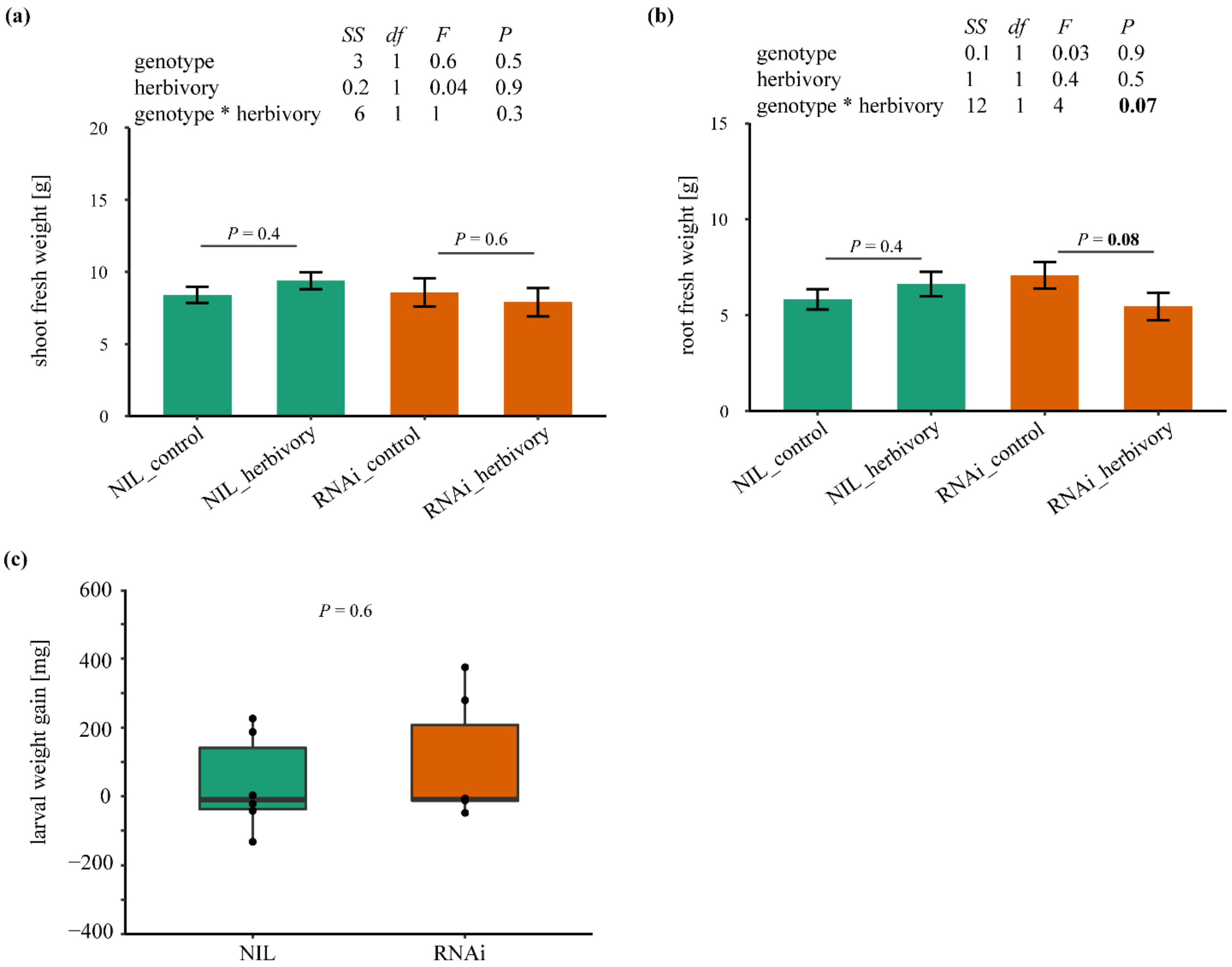


**Fig. S5** Resistance of rubber-deficient *Tk*CPTL1-RNAi and normal rubber content NIL plants of the A line to *M. melolontha* herbivory when grown on natural field soil. **(a)** Shoot and **(b)** root fresh weight after 14 days of herbivory or wounding. Compared to plant line B (Fig. 1), the pattern tended to be similar albeit not significant for root biomass (interaction plant genotype * herbivory, *P*=0.07, linear model). *P*-values of linear models are displayed. *P*-values of pairwise comparisons refer to FDR corrected values upon extraction with the function emmeans. SS=sum squares, df=degrees of freedom. **(c)** Larval weight gain was unaffected by genotype. *P*-values refer to Wilcoxon signed-rank tests. N=6.

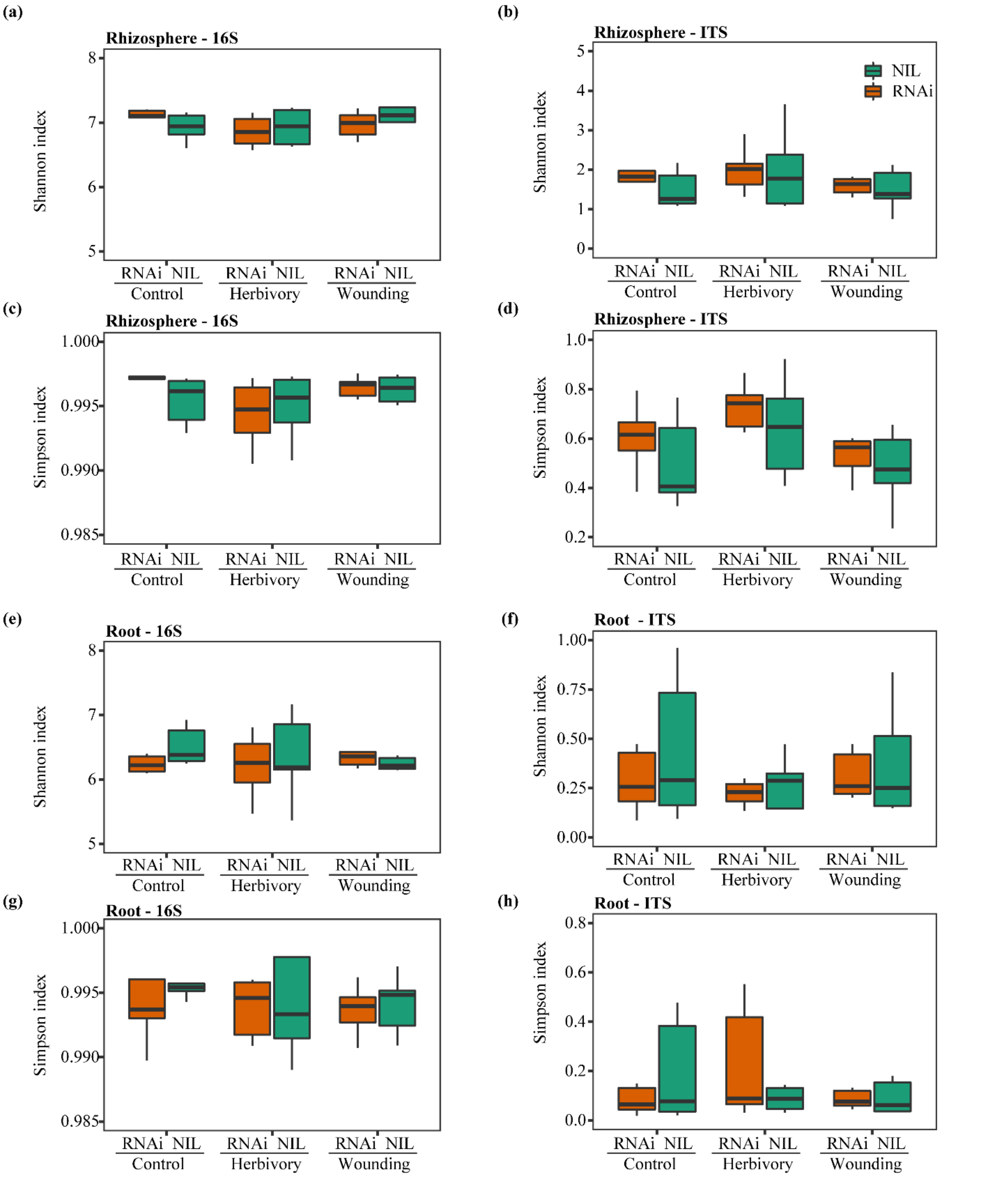


**Fig. S6** The microbial diversity was neither affected by silencing *cis*-1,4-polyisoprene biosynthesis nor the damage silencing treatment (wounding, herbivory) in the **(a-d)** rhizosphere and **(e-h)** roots based on Shannon and Simpson indexes when *T. koksaghyz* was grown on natural field soil. Statistical analysis is found in Supporting Information Table S1 – S4). N=6.

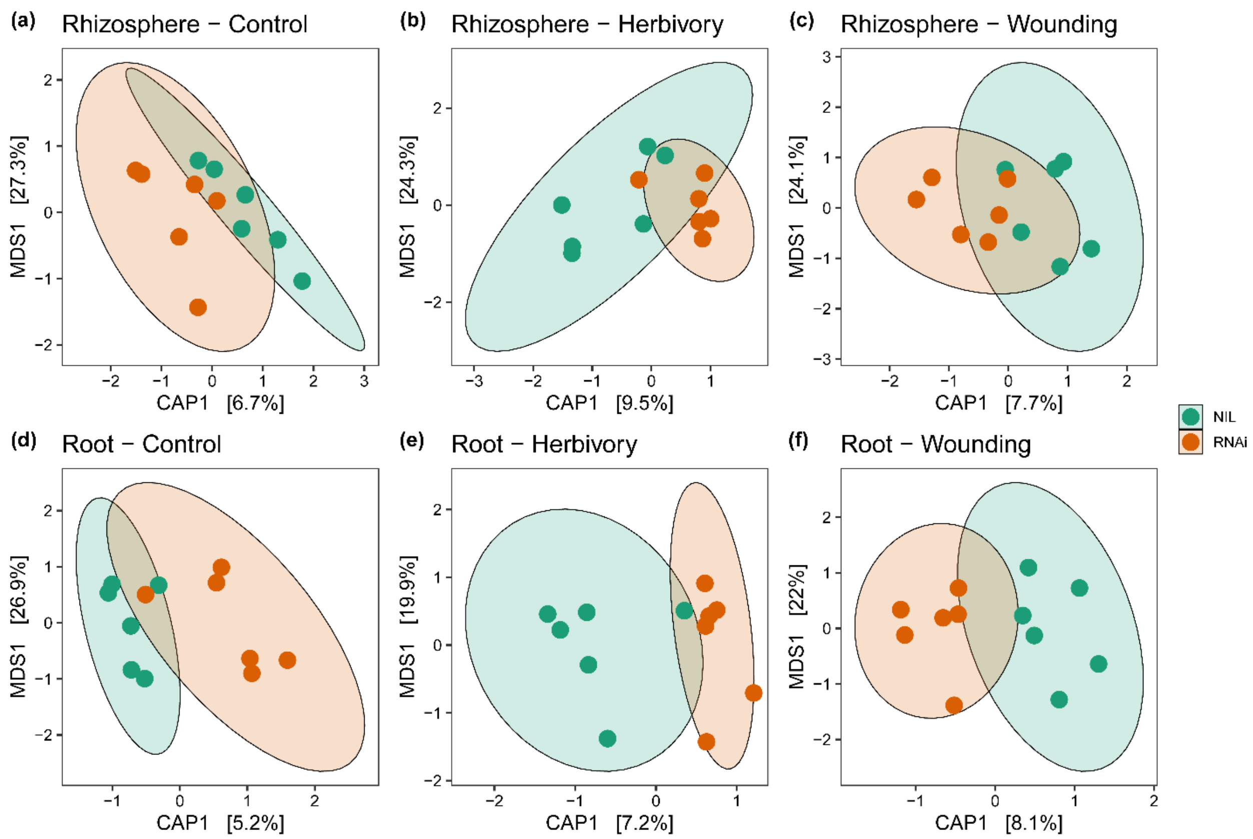


**Fig. S7** Canonical analysis of principal (CAP) coordinates ordination (Bray–Curtis distance matrix) on bacterial metabarcoding in the *T. koksaghyz* **(a, b, c)** rhizosphere and **(d, e, f)** roots, reporting the effect of genotype (NIL, *Tk*CPTL1-RNAi, different colors) under (**a, d**) control (no herbivory), **(b, e)** herbivory and **(c, f)** mechanical wounding. Percentages in parentheses report the variance explained by the respective axis. N=6.

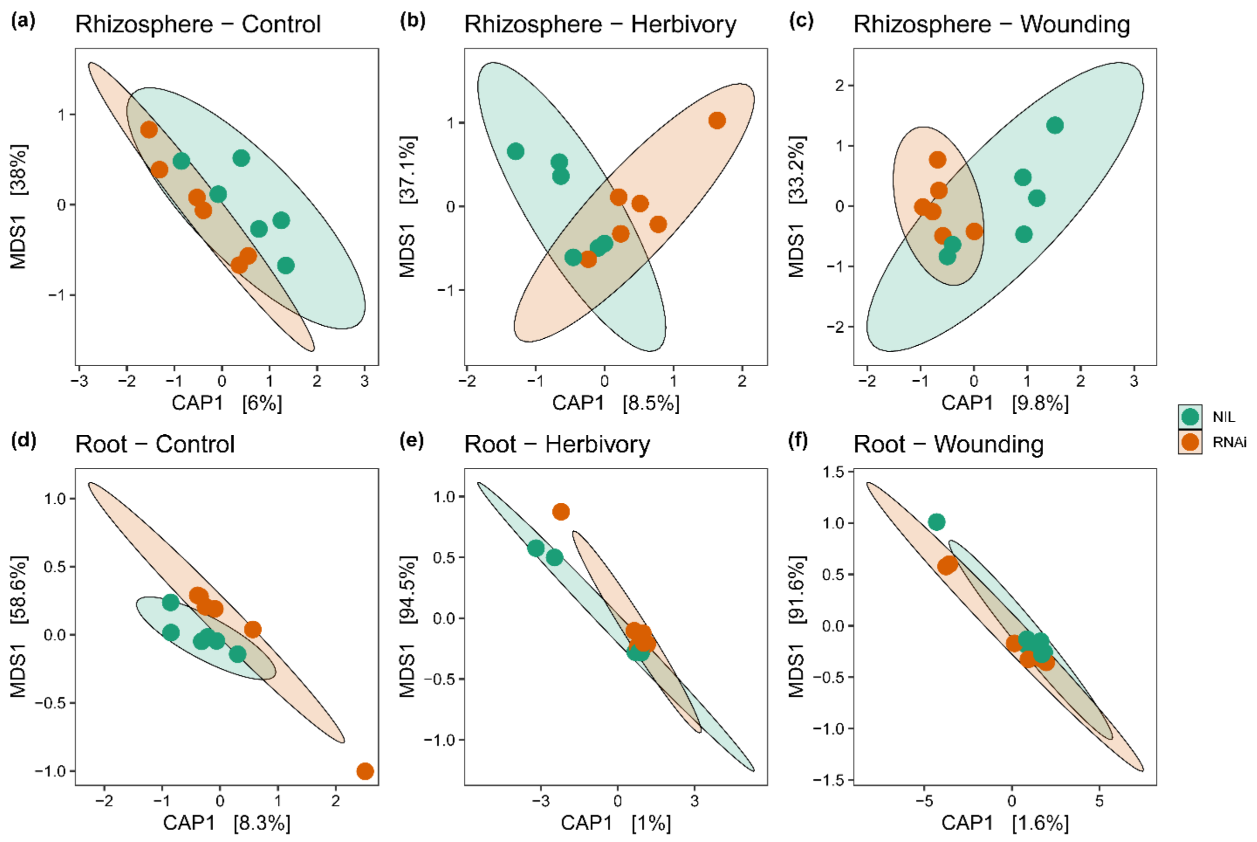


**Fig. S8** Canonical analysis of principal (CAP) coordinates ordination (Bray–Curtis distance matrix) on fungal metabarcoding in the *T. koksaghyz* **(a, b, c)** rhizosphere and **(d, e, f)** roots, reporting the effect of genotype (NIL, *Tk*CPTL1-RNAi, different colors) under **(a, d)** control (no herbivory), (**b, e**) herbivory and **(c, f)** mechanical wounding treatment. Percentages in parentheses report the variance explained by the respective axis. N=6.

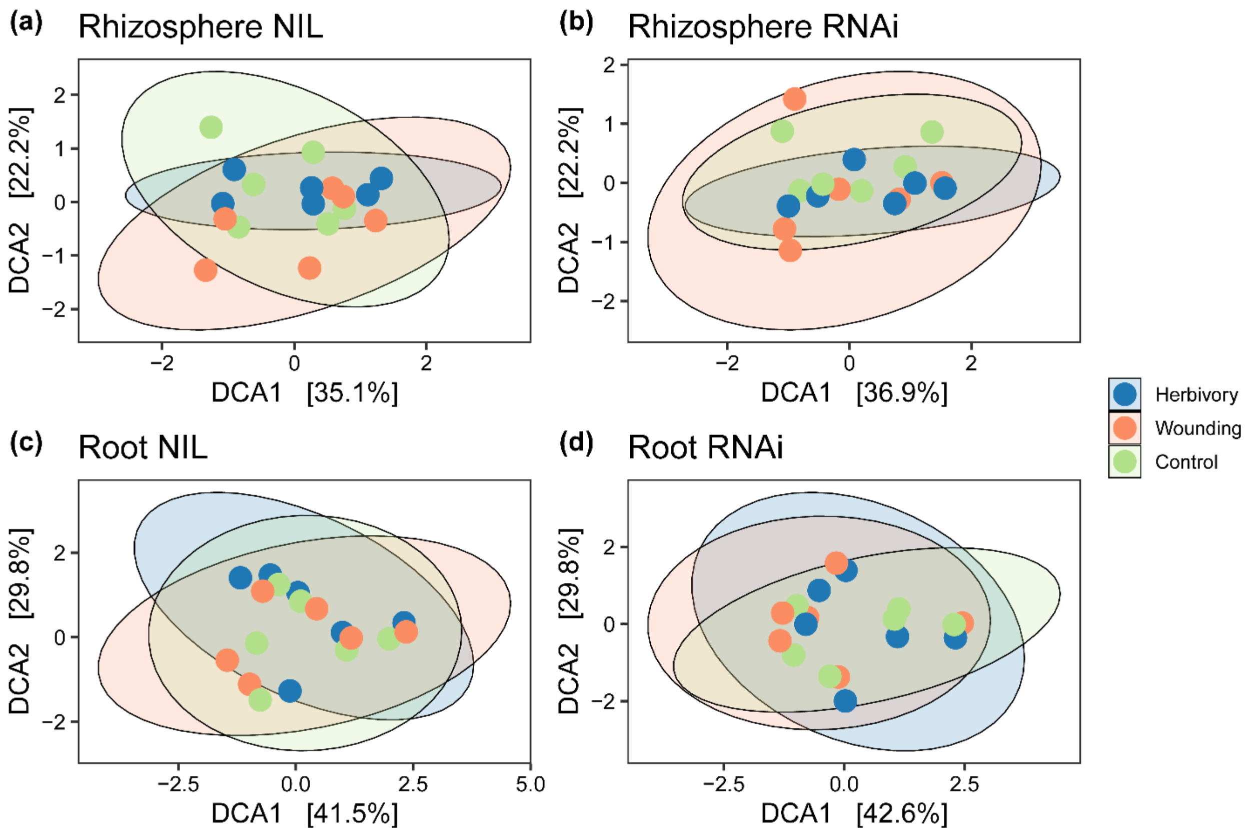


**Fig. S9** Detrended correspondence analysis (DCA) ordination (Bray–Curtis distance matrix) of **(a, b)** rhizosphere and **(c, d)** root in NIL and *Tk*CPTL1-RNAi samples for the bacterial community. For each panel we highlight the response of bacterial communities (16S) to different treatments (herbivory, wounding and control, different colors). Percentages in parentheses report the variance explained by the respective axis. N=6.

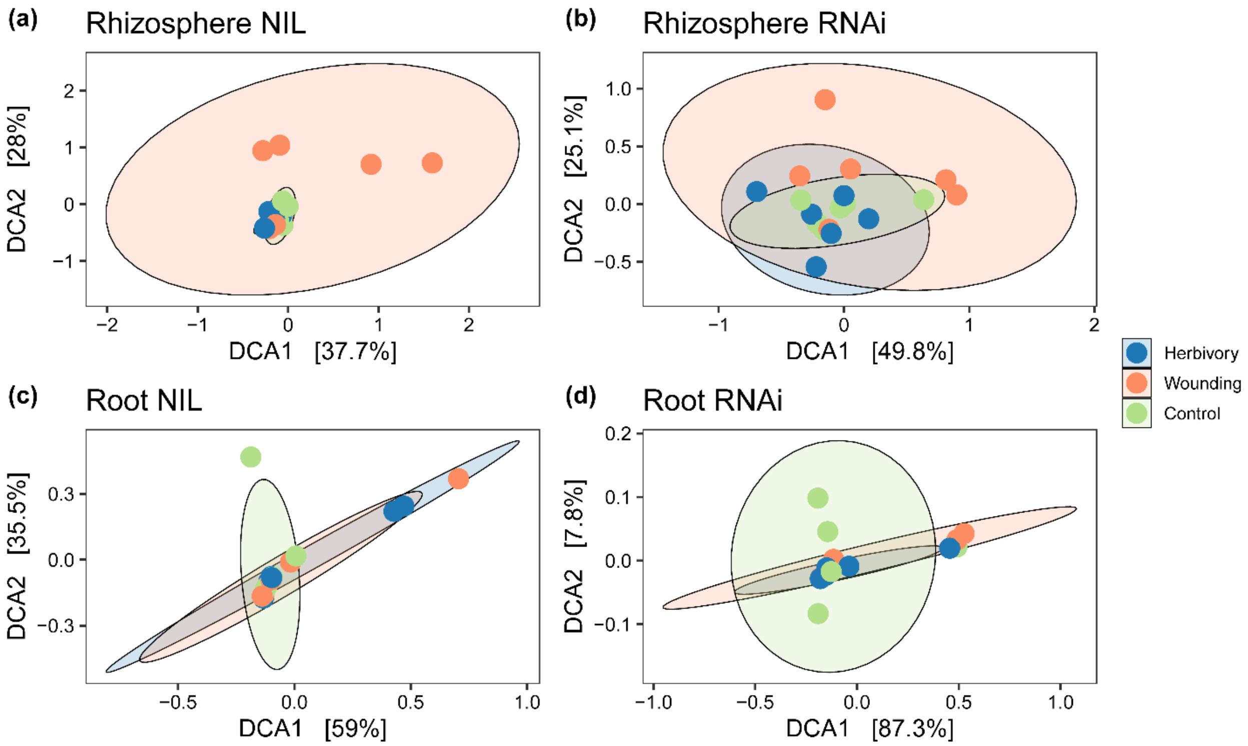


**Fig. S10** Detrended correspondence analysis (DCA) ordination (Bray–Curtis distance matrix) of **(a, b)** rhizosphere and **(c, d)** root in NIL and *Tk*CPTL1-RNAi *T. koksaghyz* plants for the fungal community. For each panel we highlight the response of fungal communities (ITS) to different treatments (herbivory, wounding and control, different colors). Percentages in parentheses report the variance explained by the respective axis. N=6.

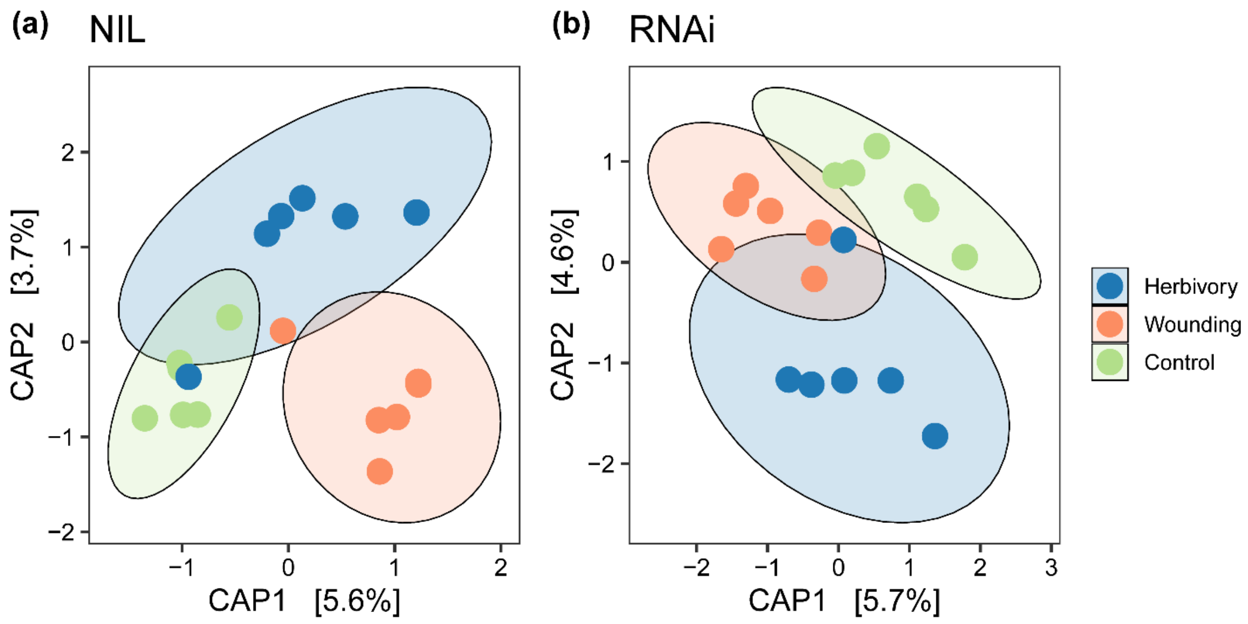


**Fig. S11** Canonical analysis of principal (CAP) coordinates ordination (Bray–Curtis distance matrix) on microbial gene content from rhizosphere samples, reporting the effect of treatment (herbivory, wounding and control, different colors) for both **(a)** NIL and **(b)** *Tk*CPTL1-RNAi genotypes. Percentages in parentheses report the variance explained by the respective axis. N=6.

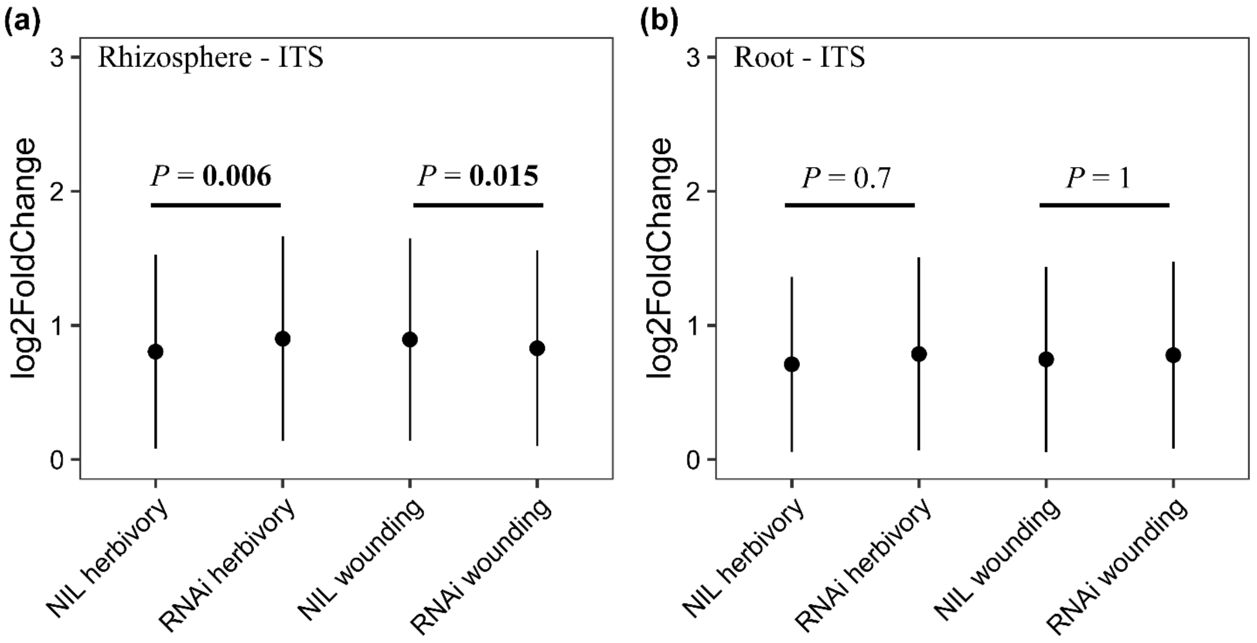


**Fig. S12** Magnitude of changes in abundance for each fungal OTU (absolute log2 fold changes). For each treatment, we investigated the response of single OTUs to the treatment compared with the respective control. Comparisons were tested using a linear mixed-effects model for **(a)** rhizosphere ($\chi2$=30.84, df=3, $P$<0.001) and **(b)** root samples ($\chi2$=3.71, df=3, $P$=0.29). Contrasts were extracted using the function emmeans (FDR corrected) and are also reported in the Supporting Information Table S5. Error bars = SE. N=6.

**Table S1** Microbial diversity analysis based on Shannon index (model coefficients).

| | Effect | Rhizosphere | | Root | |
|---|---|---|---|---|---|
| | | **Estimate** | **95% CI** | **Estimate** | **95% CI** |
| **16S** | Intercept | 7.077 | (6.926 – 7.235) | 6.237 | (5.954 – 6.513) |
| | *Plant genotype* **NIL** | - 0.137 | (-0.354 – 0.079) | 0.231 | (-0.144 – 0.617) |
| | *Treatment* **Control** | - 0.109 | (-0.329 – 0.103) | 0.143 | (-0.246 – 0.528) |
| | *Treatment* **Wounding** | - 0.194 | (-0.418 – 0.027) | -0.002 | (-0.376 – 0.369) |
| | *Plant genotype* **NIL** * *Treatment* **Control** | 0.148 | (-0.157 – 0.426) | -0.475 | (-1.016 – 0.073) |
| | *Plant genotype* **NIL** * *Treatment* **Wounding** | 0.194 | (-0.106 – 0.513) | -0.119 | (-0.649 – 0.411) |
| **ITS** | Intercept | 1.848 | (1.505 – 2.206) | 0.370 | (0.175 – 0.571) |
| | *Plant genotype* **NIL** | -0.319 | (-0.828 – 0.170) | 0.065 | (-0.205 – 0.326) |
| | *Treatment* **Control** | -0.231 | (-0.724 – 0.260) | 0.028 | (-0.257 – 0.308) |
| | *Treatment* **Wounding** | 0.098 | (-0.387 – 0.578) | 0.120 | (-0.153 – 0.397) |
| | *Plant genotype* **NIL** * *Treatment* **Control** | 0.238 | (-0.482 – 0.949) | -0.084 | (-0.474 – 0.322) |
| | *Plant genotype* **NIL** * *Treatment* **Wounding** | 0.300 | (-0.396 – 0.979) | -0.148 | (-0.537 – 0.245) |

Model coefficients from the linear model fit to the Shannon diversity index using Bayesian Hamiltonian Markov chain Monte Carlo, for both 16S and ITS datasets obtained from rhizosphere and root. All estimates resided within the 95% confidence interval (CI).

**Table S2** Microbial diversity analysis based on Simpson index (model coefficients).

| | Effect | Rhizosphere | | Root | |
|---|---|---|---|---|---|
| | | **Estimate** | **95% CI** | **Estimate** | **95% CI** |
| **16S** | Intercept | 0.997 | (0.995 – 0.998) | 0.989 | (0.979 – 1.000) |
| | *Plant genotype* **NIL** | -0.001 | (-0.003 – 0.001) | 0.005 | (-0.009 – 0.020) |
| | *Treatment* **Control** | -0.001 | (-0.003 – 0.001) | 0.004 | (-0.012 – 0.019) |
| | *Treatment* **Wounding** | -0.002 | (-0.004 – 0.001) | 0.001 | (-0.015 – 0.016) |
| | *Plant genotype* **NIL** * *Treatment* **Control** | 0.001 | (-0.002 – 0.004) | -0.018 | (-0.040 – 0.02) |
| | *Plant genotype* **NIL** * *Treatment* **Wounding** | 0.002 | (-0.001 – 0.004) | -0.006 | (-0.028 – 0.014) |
| **ITS** | Intercept | 0.599 | (0.502 – 0.697) | 0.146 | (0.028 – 0.263) |
| | *Plant genotype* **NIL** | -0.086 | (-0.223 – 0.046) | 0.043 | (-0.114 – 0.204) |
| | *Treatment* **Control** | -0.061 | (-0.196 – 0.073) | 0.006 | (-0.157 – 0.165) |
| | *Treatment* **Wounding** | 0.072 | (-0.061 – 0.202) | 0.065 | (-0.090 – 0.225) |
| | *Plant genotype* **NIL** * *Treatment* **Control** | 0.043 | (-0.152 – 0.229) | -0.060 | (-0.283 – 0.164) |
| | *Plant genotype* **NIL** * *Treatment* **Wounding** | 0.044 | (-0.148 – 0.234) | -0.097 | (-0.318 – 0.126) |

Model coefficients from the linear model fit to the Simpson diversity index using Bayesian Hamiltonian Markov chain Monte Carlo, for both 16S and ITS datasets obtained from rhizosphere and root. All estimates resided within the 95% confidence interval (CI).

**Table S3** Microbial diversity analysis based on Shannon index (pairwise comparisons).

| Pairwise comparison | Rhizosphere | | | | Root | | | |
|---|---|---|---|---|---|---|---|---|
| | 16S | | ITS | | 16S | | ITS | |
| | Estimate | pMCMC | Estimate | pMCMC | Estimate | pMCMC | Estimate | pMCMC |
| **NIL Herb. - NIL Ctr** | -0.0387 | 0.7735 | -0.0071 | 0.9870 | 0.3320 | 0.1420 | 0.0555 | 0.7445 |
| **NIL Wound. - NIL Ctr** | -0.0385 | 0.7900 | 0.3903 | 0.1800 | 0.2102 | 0.3600 | 0.0273 | 0.8885 |
| **NIL Herb. – NIL Wound.** | -0.0002 | 0.9970 | -0.3975 | 0.1790 | 0.1217 | 0.5740 | 0.0282 | 0.8640 |
| **TR Herb. – TR Ctr** | 0.1091 | 0.4130 | 0.2311 | 0.4270 | -0.1427 | 0.5350 | -0.0281 | 0.8735 |
| **TR Wound. - TR Ctr** | -0.0846 | 0.5225 | 0.3292 | 0.2780 | -0.1452 | 0.5285 | 0.0919 | 0.5745 |
| **TR Herb. – TR Wound.** | 0.1937 | 0.1440 | -0.0981 | 0.7305 | 0.0024 | 0.9930 | -0.12001 | 0.4835 |
| **NIL Ctr – TR Ctr** | 0.0111 | 0.9230 | -0.0806 | 0.7915 | -0.2438 | 0.2870 | -0.0182 | 0.9330 |
| **NIL Herb. - TR Herb.** | -0.1366 | 0.3000 | -0.3191 | 0.2905 | 0.2309 | 0.3100 | 0.0653 | 0.6875 |
| **NIL Wound. - TR Wound.** | 0.0573 | 0.6855 | -0.0195 | 0.9470 | 0.1116 | 0.6135 | -0.0828 | 0.6300 |

Pairwise comparisons obtained from fitting a linear mixed-effect model using Bayesian Hamiltonian Markov chain Monte Carlo on Shannon index for both 16S and ITS in rhizosphere and root samples. Herb=Herbivory treatment, Wound=Wounding treatment, Ctr=control treatment, TR=treatment. None of the pairwise comparisons showed significant differences.

**Table S4** Microbial diversity analysis (pairwise comparisons).

| Pairwise comparison | Rhizosphere | | | | Root | | | |
|---|---|---|---|---|---|---|---|---|
| | 16S | | ITS | | 16S | | ITS | |
| | Estimate | pMCMC | Estimate | pMCMC | Estimate | pMCMC | Estimate | pMCMC |
| **NIL Herb. - NIL Ctr** | -8.6e-05 | 0.9285 | 0.0185 | 0.8085 | 0.0145 | 0.1075 | 0.0547 | 0.5495 |
| **NIL Wound. - NIL Ctr** | -4.6e-04 | 0.6990 | 0.1347 | 0.1065 | 0.0088 | 0.3220 | 0.0224 | 0.8245 |
| **NIL Herb. - NIL Wound.** | 3.8e-04 | 0.7625 | -0.1161 | 0.1415 | 0.0056 | 0.5155 | 0.0323 | 0.7250 |
| **TR Herb. – TR Ctr** | 1.1e-03 | 0.3440 | 0.0613 | 0.4620 | -0.0038 | 0.6735 | -0.0056 | 0.9345 |
| **TR Wound. - TR Ctr** | -9.7e-04 | 0.3955 | 0.1333 | 0.0970 | -0.00315 | 0.7290 | 0.0592 | 0.5270 |
| **TR Herb. – TR Wound.** | 2.04e-03 | 0.0800 | -0.0721 | 0.3775 | -0.0006 | 0.9245 | -0.0648 | 0.5065 |
| **NIL Ctr – TR Ctrl** | -4.81e-05 | 0.9835 | -0.0428 | 0.5770 | -0.01306 | 0.1465 | -0.0177 | 0.8475 |
| **NIL Herb. - TR Herb.** | -1.2e-03 | 0.3030 | -0.0855 | 0.2975 | 0.00531 | 0.5700 | 0.0427 | 0.6585 |
| **NIL Wound. - TR Wound.** | 4.6e-04 | 0.7015 | -0.0414 | 0.6110 | -0.0011 | 0.9175 | -0.0544 | 0.5765 |

Pairwise comparisons obtained from fitting a linear mixed-effect model using Bayesian Hamiltonian Markov chain Monte Carlo on Simpson index for both 16S and ITS in rhizosphere and root samples. Herb=Herbivory treatment, Wound=Wounding treatment, Ctr=control treatment, TR=treatment. None of the pairwise comparisons showed significant differences.

**Table S5** Magnitude of changes in abundance for each bacterial and fungal OTU (absolute log2 fold changes).

| **Contrast** | **Bacterial OTU** | | | | **Fungal OTU** | | | |
|---|---|---|---|---|---|---|---|---|
| | **Rhizosphere** | | **Root** | | **Rhizosphere** | | **Root** | |
| | **Estimate** | ***P*** | **Estimate** | ***P*** | **Estimate** | ***P*** | **Estimate** | ***P*** |
| **RNAi Herbivory - RNAi Wounding** | 0.0277 | **0.0368** | -0.1081 | **<.0001** | 0.0346 | 0.8599 | -0.03056 | 0.9405 |
| **RNAi Herbivory - NIL Herbivory** | 0.3959 | **<.0001** | 0.0533 | **<.0001** | 0.1447 | **0.0064** | 0.06368 | 0.6517 |
| **RNAi Herbivory - NIL Wounding** | 0.0562 | **<.0001** | 0.0131 | 0.6076 | -0.0985 | 0.1200 | -0.02241 | 0.9770 |
| **RNAi Wounding - NIL Herbivory** | 0.3682 | **<.0001** | 0.1615 | **<.0001** | 0.1101 | 0.0645 | 0.09423 | 0.3144 |
| **RNAi Wounding - NIL Wounding** | 0.0285 | **0.0314** | 0.1212 | **<.0001** | -0.1332 | **0.0148** | 0.00815 | 0.9988 |
| **NIL Herbivory - NIL Wounding** | -0.3396 | **<.0001** | -0.0403 | **0.0006** | -0.2432 | **<.0001** | -0.08609 | 0.3781 |

For each treatment, we investigated the response of single OTUs to the treatment in comparison with the control of the respective genotype. Comparisons were tested using a linear mixed-effects model for bacterial rhizosphere samples ($\chi2=1900$, df=3, $P<0.001$), bacterial root samples ($\chi2=250.64$, df=3, $P<0.001$), fungal rhizosphere samples ($\chi2=30.84$, df=3, $P<0.001$) and fungal root samples ($\chi2=3.71$, df=3, $P=0.29$). Contrasts for OTUs were extracted using the function emmeans (FDR corrected). See also Fig. 3b and Fig. S12.

**Table S6** Number of observed OTUs that are differentially abundant between each treatment group and the respective control in the rhizosphere 16S and ITS dataset.

| Group | Rhizosphere 16S | | | | Rhizosphere ITS | | | |
|---|---|---|---|---|---|---|---|---|
| | Observed OTUs | Random OTUs | $\chi^2$ | *P* | Observed OTUs | Random OTUs | $\chi^2$ | *P* |
| **NIL Herbivory** | 0 | 33 | 33.00 | < 0.001 | 0 | 15 | 15.00 | < 0.001 |
| **NIL Wounding** | 117 | 9 | 92.57 | < 0.001 | 4 | 11 | 3.26 | 0.071 |
| **RNAi Herbivory** | 11 | 40 | 16.49 | < 0.001 | 0 | 18 | 18.00 | < 0.001 |
| **RNAi Wounding** | 5 | 38 | 25.32 | < 0.001 | 0 | 10 | 10.00 | 0.001 |

The random number of OTUs was generated by randomizing (1000 iterations) the counts within the same OTU table but keeping constant the number of reads per sample. Differences between the number of observed OTUs and the number of random OTUs were assessed using a $\chi2$ test.

**Table S7** Number of observed OTUs that are differentially abundant between each treatment group and the respective control in the rhizosphere 16S dataset.

| Plant genotype | Treatment | Direction | Genus | # of OTUs |
|---|---|---|---|---|
| **NIL** | **Herbivory** | **Up** | - | - |
| | | **Down** | - | - |
| | **Wounding** | **Up** | *Pseudonocardia* | 1 |
| | | | Unidentified | 1 |
| | | **Down** | *Acidibacter* | 1 |
| | | | *Aeromicrobium* | 1 |
| | | | *Agromyces* | 1 |
| | | | *Alicyclobacillus* | 1 |
| | | | *Altererythrobacter* | 1 |
| | | | *Arthrobacter* | 2 |
| | | | *BD1-7_clade* | 1 |
| | | | *Bacillus* | 3 |
| | | | *Blastococcus* | 1 |
| | | | *Bryobacter* | 1 |
| | | | *Candidatus_Alysiosphaera* | 1 |
| | | | *Candidatus_Nitrososphaera* | 1 |
| | | | *Clostridium_sensu_stricto_13* | 1 |
| | | | *Defluviicoccus* | 2 |

| | | |
|---|---|---|
| | *Devosia* | 1 |
| | *Fictibacillus* | 1 |
| | *Flavobacterium* | 1 |
| | *Gaiella* | 2 |
| | *Haliangium* | 1 |
| | *Halomonas* | 1 |
| | *Legionella* | 1 |
| | *Lysobacter* | 1 |
| | *Mesorhizobium* | 2 |
| | *Methylibium* | 1 |
| | *Mycobacterium* | 1 |
| | *Nitrosospira* | 1 |
| | *Nocardioides* | 2 |
| | *Nocardioides* | 1 |
| | *Opitutus* | 1 |
| | *Paenibacillus* | 3 |
| | *Pedomicrobium* | 1 |
| | *Phenylobacterium* | 1 |
| | *Promicromonospora* | 1 |
| | *Pseudolabrys* | 1 |

| | | | | |
|---|---|---|---|---|
| | | | *Pseudomonas* | 2 |
| | | | *Pseudoxanthomonas* | 1 |
| | | | *Rhizobium* | 3 |
| | | | *Rhizocola* | 1 |
| | | | *Rubrobacter* | 1 |
| | | | *Shinella* | 1 |
| | | | *Solirubrobacter* | 1 |
| | | | *Sorangium* | 1 |
| | | | *Sphingobium* | 2 |
| | | | *Sphingomonas* | 3 |
| | | | *Sphingopyxis* | 1 |
| | | | *Stenotrophomonas* | 1 |
| | | | *Streptomyces* | 2 |
| | | | *Thermobacillus* | 1 |
| | | | *Tumebacillus* | 1 |
| | | | *Turicibacter* | 1 |
| | | | Unidentified | 55 |
| **RNAi** | **Herbivory** | **Up** | *Alicyclobacillus* | 1 |
| | | | *Devosia* | 1 |
| | | | *Emticicia* | 1 |

| | | | |
|---|---|---|---|
| | | *Microbacterium* | 1 |
| | | *Nocardioides* | 1 |
| | **Down** | *Pseudomonas* | 1 |
| | | Unidentified | 1 |
| **Wounding** | **Up** | - | - |
| | **Down** | *Pseudomonas* | 1 |

For each treatment group and direction ('up' refers to increase compared to control, 'down' refers to a decrease compared to control) we report the number of OTUs for each bacteria genus.

**Table S8** Number of observed OTUs that are differentially abundant between each treatment group and the respective control in the rhizosphere ITS dataset.

| | Treatment | Direction | Genus | # of OTUs |
|---|---|---|---|---|
| NIL | Herbivory | Up | - | - |
| | | Down | - | - |
| | Wounding | Up | *Bjerkandera* | 1 |
| | | | *Gallinipes* | 1 |
| | | | *Toxicocladosporium* | 1 |
| | | | *Unidentified* | 1 |
| | | Down | - | - |
| RNAi | Herbivory | Up | - | - |
| | | Down | - | - |
| | Wounding | Up | - | - |
| | | Down | - | - |

For each treatment group and direction (‘up’ refers to increase compared to control, ‘down’ refers to a decrease compared to control) we report the number of OTUs for each fungal genus.

**Table S9** Number of observed OTUs that are differentially abundant between each treatment group and the respective control in the root 16S and ITS dataset.

| | Root 16S | | | | Root ITS | | | |
|---|---|---|---|---|---|---|---|---|
| **Group** | **Observed OTUs** | **Random OTUs** | $\chi^2$ | ***P*** | **Observed OTUs** | **Random OTUs** | $\chi^2$ | ***P*** |
| **NIL Herbivory** | 0 | 0 | *NA* | *NA* | 0 | 1 | 1 | 0.3 |
| **NIL Wounding** | 0 | 0 | *NA* | *NA* | 0 | 1 | 1 | 0.3 |
| **RNAi Herbivory** | 1 | 0 | 1 | 0.3 | 0 | 1 | 1 | 0.3 |
| **RNAi Wounding** | 0 | 3 | 3 | 0.08 | 0 | 2 | 2 | 0.3 |

The random number of OTUs was generated by randomizing (1000 iterations) the counts within the same OTU table but keeping constant the number of reads per sample. Differences between the number of observed OTUs and the number of random OTUs were assessed using a $\chi 2$ test.

**Table S10** Number of observed OTUs that are differentially abundant between each treatment group and the respective control in the root 16S dataset.

| Plant genotype | Treatment | Direction | Genus | # of OTUs |
|---|---|---|---|---|
| **NIL** | **Herbivory** | **Up** | - | - |
| | | **Down** | - | - |
| | **Wounding** | **Up** | - | - |
| | | **Down** | - | - |
| **RNAi** | **Herbivory** | **Up** | *Arthrobacter* | 1 |
| | | **Down** | - | - |
| | **Wounding** | **Up** | - | - |
| | | **Down** | - | - |

For each treatment group and direction ('up' refers to increase compared to control, 'down' refers to a decrease compared to control) we report the number of OTUs for each bacterial genus.

**Table S11** Relative abundance of bacterial genus in plant rhizosphere.

| Plant genotype | Treatment | Genus | F | *p* | Rel. abundance treatment (%) | Rel. abundance control (%) |
|---|---|---|---|---|---|---|
| **NIL** | **herbivory** | *Arthrobacter* | 9.13 | 0.012 | **8.97** | 3.70 |
| **NIL** | **wounding** | *Arthrobacter* | 22.74 | <0.001 | **9.07** | 3.70 |
| | | *Rhizobium* | 8.87 | 0.013 | 4.93 | **7.67** |
| **RNAi** | **herbivory** | *Mesorhizobium* | 8.47 | 0.015 | **1.22** | 0.83 |
| | | *Mycobacterium* | 13.67 | 0.004 | **3.62** | 1.91 |
| **RNAi** | **wounding** | *Devosia* | 7.96 | 0.028 | 1.56 | **2.14** |
| | | *Microbacterium* | 8.28 | 0.026 | **1.38** | 0.80 |
| | | *Mycobacterium* | 11.58 | 0.006 | **2.44** | 1.91 |
| | | *Novosphingobium* | 13.65 | 0.004 | **3.58** | 2.05 |
| | | *Opitutus* | 5.21 | 0.045 | 1.01 | **1.72** |
| | | *Pseudomonas* | 6.21 | 0.03 | **11.8** | 6.66 |

Comparisons were performed between treatments and their relative control group using a linear model. Values in bold indicate groups with higher abundance.

**Table S12** Relative abundance of bacterial genus in plant roots.

| Plant genotype | Treatment | Genus | F | *p* | Rel. abundance treatment (%) | Rel. abundance control (%) |
|---|---|---|---|---|---|---|
| **NIL** | **herbivory** | *Bacillus* | 5.06 | 0.048 | **7.72** | 3.42 |
| | | *Cellvibrio* | 4.97 | 0.049 | 1.40 | **2.07** |
| | | *Mycobacterium* | 6.20 | 0.031 | **3.99** | 1.94 |
| **NIL** | **wounding** | *Cellvibrio* | 20.71 | 0.001 | 1.32 | **2.07** |
| | | *Novosphingobium* | 8.23 | 0.016 | 1.63 | **2.55** |
| | | *Rhizobium* | 18.01 | 0.001 | 5.51 | **8.48** |
| | | *Sphingobium* | 5.45 | 0.041 | 5.23 | **7.88** |
| **RNAi** | **herbivory** | *Bacillus* | 7.51 | 0.021 | 3.49 | **5.48** |
| | | *Paenibacillus* | 6.11 | 0.033 | 0.99 | **1.49** |
| **RNAi** | **wounding** | *Bacillus* | 13.35 | 0.004 | 2.90 | **5.48** |
| | | *Paenibacillus* | 8.91 | 0.013 | 0.89 | **1.49** |
| | | *Pseudomonas* | 6.41 | 0.029 | 19.53 | **10.55** |

Comparisons were performed between treatments and their relative control group using a linear model. Values in bold indicate groups with higher abundance.

**Table S13** Functional analysis displaying the number of observed genes that are differentially abundant between each treatment group and the respective control in the rhizosphere.

| Group | Observed genes | Random genes | $\chi^2$ | *P* |
|---|---|---|---|---|
| **NIL herbivory** | 1079 | 37 | 972.91 | < 0.001 |
| **NIL wounding** | 1179 | 27 | 1100.41 | < 0.001 |
| **RNAi herbivory** | 496 | 65 | 331.12 | < 0.001 |
| **RNAi wounding** | 67 | 91 | 3.64 | 0.056 |

The random number of OTUs was generated by randomizing (1000 iterations) the counts within the same OTU table but keeping constant the number of reads per sample. See also Figure 4b.

**Table S14** List of genes and corresponding group stating a higher abundance.

| Gene (NIL-Herbivory vs NIL-Control) | Group with higher abundance |
|---|---|
| 1-deoxy-D-xylulose-5-phosphate synthase | NIL–Herbivory |
| 1,4-alpha-glucan branching enzyme GlgB | NIL–Herbivory |
| 2-aminomuconic 6-semialdehyde dehydrogenase | NIL–Herbivory |
| 3-ketoacyl-CoA thiolase | NIL–Herbivory |
| 3-methylmercaptopropionyl-CoA dehydrogenase | NIL–Herbivory |
| 3-phenylpropionate-dihydrodiol/cinnamic acid-dihydrodiol dehydrogenase | NIL–Herbivory |
| 30S ribosomal protein S1 | NIL–Herbivory |
| 4-alpha-glucanotransferase | NIL–Herbivory |
| 5-dehydro-2-deoxygluconokinase | NIL–Herbivory |
| Acetyl-coenzyme A synthetase | NIL–Herbivory |
| Adaptive-response sensory-kinase SasA | NIL–Herbivory |
| Adenosylhomocysteinase | NIL–Herbivory |
| Adenosylmethionine-8-amino-7-oxononanoate aminotransferase | NIL–Herbivory |
| Aerobic C4-dicarboxylate transport protein | NIL–Herbivory |
| Alanine--tRNA ligase | NIL–Herbivory |
| Alpha-galactosidase | NIL–Herbivory |
| Alpha-xylosidase | NIL–Herbivory |
| Amidophosphoribosyltransferase | NIL–Herbivory |
| Arginine biosynthesis bifunctional protein ArgJ | NIL–Herbivory |
| Aspartate--tRNA(Asp/Asn) ligase | NIL–Herbivory |
| Aspartyl/glutamyl-tRNA(Asn/Gln) amidotransferase subunit B | NIL–Herbivory |
| ATP-dependent DNA helicase UvrD1 | NIL–Herbivory |
| ATP-dependent RNA helicase DeaD | NIL–Herbivory |
| ATP-dependent RNA helicase SrmB | NIL–Herbivory |
| Beta-galactosidase | NIL–Herbivory |
| Beta-hexosaminidase | NIL–Herbivory |
| Beta-N-acetylglucosaminidase/beta-glucosidase | NIL–Herbivory |
| Beta-xylosidase | NIL–Herbivory |
| Bifunctional protein PaaZ | NIL–Herbivory |
| Biosynthetic peptidoglycan transglycosylase | NIL–Herbivory |
| Calcium-transporting ATPase 1 | NIL–Herbivory |
| Cardiolipin synthase A | NIL–Herbivory |
| Chaperone protein ClpB | NIL–Herbivory |
| Chaperone protein DnaK | NIL–Herbivory |
| Chromosome partition protein Smc | NIL–Herbivory |
| Cytochrome bd ubiquinol oxidase subunit 1 | NIL–Herbivory |
| D-xylose-proton symporter | NIL–Herbivory |
| Dihydroxy-acid dehydratase | NIL–Herbivory |

| | |
|---|---|
| Dipeptidyl aminopeptidase BI | NIL–Herbivory |
| Dipeptidyl-peptidase 5 | NIL–Herbivory |
| DNA gyrase subunit B | NIL–Herbivory |
| DNA helicase II | NIL–Herbivory |
| DNA polymerase I | NIL–Herbivory |
| DNA polymerase III subunit alpha | NIL–Herbivory |
| DNA-directed RNA polymerase subunit beta JHCEEJLM_17894 | NIL–Herbivory |
| Elongation factor 4 | NIL–Herbivory |
| Energy-dependent translational throttle protein EttA | NIL–Herbivory |
| Enolase | NIL–Herbivory |
| FAD-containing monooxygenase EthA | NIL–Herbivory |
| Fosfomycin resistance protein AbaF | NIL–Herbivory |
| Glucose-6-phosphate 1-dehydrogenase | NIL–Herbivory |
| GTP-binding protein TypA/BipA | NIL–Herbivory |
| High-affinity choline transport protein | NIL–Herbivory |
| HTH-type transcriptional regulator MalT | NIL–Herbivory |
| hypothetical protein | NIL–Herbivory |
| Inosine-5-monophosphate dehydrogenase CKCIAONA_26804 | NIL–Herbivory |
| Inositol 2-dehydrogenase/D-chiro-inositol 3-dehydrogenase | NIL–Herbivory |
| IS3 family transposase ISStma9 | NIL–Herbivory |
| Isocitrate dehydrogenase [NADP] | NIL–Herbivory |
| Isoleucine--tRNA ligase | NIL–Herbivory |
| Isoquinoline 1-oxidoreductase subunit beta | NIL–Herbivory |
| L-asparagine permease 1 | NIL–Herbivory |
| Leucine--tRNA ligase | NIL–Herbivory |
| Levanase | NIL–Herbivory |
| Levanbiose-producing levanase | NIL–Herbivory |
| Long-chain-fatty-acid--CoA ligase | NIL–Herbivory |
| Long-chain-fatty-acid--CoA ligase FadD15 | NIL–Herbivory |
| Low affinity potassium transport system protein kup | NIL–Herbivory |
| Malate synthase | NIL–Herbivory |
| Mannitol 2-dehydrogenase | NIL–Herbivory |
| Mannuronan C5-epimerase | NIL–Herbivory |
| Metal-pseudopaline receptor CntO | NIL–Herbivory |
| Methionine--tRNA ligase | NIL–Herbivory |
| Molybdenum import ATP-binding protein ModC | NIL–Herbivory |
| Multidrug efflux pump subunit AcrB | NIL–Herbivory |
| Multidrug resistance protein 3 | NIL–Herbivory |
| N-acetylglucosaminyldiphosphoundecaprenol N-acetyl-beta-D-mannosaminyltransferase | NIL–Herbivory |
| Na(+), Li(+), K(+)/H(+) antiporter | NIL–Herbivory |

| | |
|---|---|
| NAD(P) transhydrogenase subunit beta | NIL–Herbivory |
| NAD(P)H-quinone oxidoreductase subunit 2, chloroplastic | NIL–Herbivory |
| Neutral endopeptidase | NIL–Herbivory |
| Nitrite reductase [NAD(P)H] | NIL–Herbivory |
| O-acetyltransferase OatA | NIL–Herbivory |
| Oligopeptide-binding protein OppA | NIL–Herbivory |
| Peptidoglycan D,D-transpeptidase MrdA | NIL–Herbivory |
| Phenylacetate-coenzyme A ligase | NIL–Herbivory |
| Phenylalanine--tRNA ligase beta subunit | NIL–Herbivory |
| Phosphoenolpyruvate carboxykinase [GTP] | NIL–Herbivory |
| Phosphoenolpyruvate carboxylase | NIL–Herbivory |
| Phosphoenolpyruvate-protein phosphotransferase | NIL–Herbivory |
| Phosphomannomutase/phosphoglucomutase | NIL–Herbivory |
| Phosphoribosylformylglycinamidine synthase subunit PurL | NIL–Herbivory |
| Polyol:NADP oxidoreductase | NIL–Herbivory |
| Polyphosphate kinase | NIL–Herbivory |
| Potassium transporter KimA | NIL–Herbivory |
| Protein-methionine-sulfoxide reductase catalytic subunit MsrP | NIL–Herbivory |
| PTS system fructose-specific EIIABC component | NIL–Herbivory |
| Putative 3-oxopropanoate dehydrogenase | NIL–Herbivory |
| putative ABC transporter ATP-binding protein | NIL–Herbivory |
| putative glycine dehydrogenase (decarboxylating) | NIL–Herbivory |
| Putative tartrate transporter | NIL–Herbivory |
| Putrescine importer PuuP | NIL–Herbivory |
| Pyruvate dehydrogenase E1 component | NIL–Herbivory |
| RecBCD enzyme subunit RecB | NIL–Herbivory |
| Ribonuclease J | NIL–Herbivory |
| Ribonucleoside-diphosphate reductase subunit alpha 2 | NIL–Herbivory |
| Sarcosine oxidase subunit alpha | NIL–Herbivory |
| Sensor histidine kinase RcsC | NIL–Herbivory |
| Sensor protein KdpD | NIL–Herbivory |
| Spermidine/putrescine import ATP-binding protein PotA | NIL–Herbivory |
| Succinate--CoA ligase [ADP-forming] subunit beta | NIL–Herbivory |
| Succinate-semialdehyde dehydrogenase | NIL–Herbivory |
| Sulfite reductase [NADPH] flavoprotein alpha-component | NIL–Herbivory |
| Transcription-repair-coupling factor | NIL–Herbivory |
| Transketolase | NIL–Herbivory |
| Translation initiation factor IF-2 | NIL–Herbivory |
| Trehalase | NIL–Herbivory |
| Tryptophan 2-monooxygenase | NIL–Herbivory |
| Tyrosine recombinase XerC | NIL–Herbivory |

| | |
|---|---|
| Urocanate hydratase | NIL–Herbivory |
| UvrABC system protein A | NIL–Herbivory |
| Validamycin A dioxygenase | NIL–Herbivory |
| Valine--tRNA ligase | NIL–Herbivory |
| Vitamin B12 import ATP-binding protein BtuD | NIL–Herbivory |
| Heme transporter BhuA | NIL-Control |
| hypothetical protein | NIL-Control |
| IS256 family transposase ISEc39 | NIL-Control |

| **Gene (NIL–Wounding vs NIL-Control)** | **Group with higher abundance** |
|---|---|
| 2-amino-5-chloromuconic acid deaminase | NIL-Wounding |
| 2-aminoadipate transaminase | NIL-Wounding |
| 2-hydroxy-3-oxopropionate reductase | NIL-Wounding |
| 3-carboxy-cis,cis-muconate cycloisomerase | NIL-Wounding |
| 3-hydroxybenzoate 4-monooxygenase | NIL-Wounding |
| 3-isopropylmalate dehydrogenase | NIL-Wounding |
| 3-phenylpropionate-dihydrodiol/cinnamic acid-dihydrodiol dehydrogenase | NIL-Wounding |
| 3D-(3,5/4)-trihydroxycyclohexane-1,2-dione hydrolase | NIL-Wounding |
| 4-hydroxybenzoate 3-monooxygenase (NAD(P)H) | NIL-Wounding |
| 6-phosphogluconate dehydrogenase, NADP(+)-dependent, decarboxylating | NIL-Wounding |
| 60 kDa chaperonin 1 | NIL-Wounding |
| Acetolactate synthase large subunit IlvB1 | NIL-Wounding |
| Acetyl-coenzyme A synthetase | NIL-Wounding |
| Acyl-CoA dehydrogenase | NIL-Wounding |
| Adaptive-response sensory-kinase SasA | NIL-Wounding |
| Adenylosuccinate lyase | NIL-Wounding |
| Alanine--tRNA ligase | NIL-Wounding |
| Amino-acid carrier protein AlsT | NIL-Wounding |
| Amino-acid permease RocE | NIL-Wounding |
| Aminopeptidase N | NIL-Wounding |
| Apo-petrobactin exporter | NIL-Wounding |
| Aspartyl/glutamyl-tRNA(Asn/Gln) amidotransferase subunit B | NIL-Wounding |
| ATP synthase subunit alpha | NIL-Wounding |
| ATP-dependent 6-phosphofructokinase | NIL-Wounding |
| ATP-dependent Clp protease ATP-binding subunit ClpC1 | NIL-Wounding |
| ATP-dependent DNA helicase RecG | NIL-Wounding |
| ATP-dependent RNA helicase DeaD | NIL-Wounding |
| ATP-dependent RNA helicase RhlE | NIL-Wounding |
| ATP-dependent zinc metalloprotease FtsH | NIL-Wounding |
| Beta-xylosidase | NIL-Wounding |

| | |
|---|---|
| Bifunctional chorismate mutase/prephenate dehydratase | NIL-Wounding |
| Bifunctional protein GlmU | NIL-Wounding |
| Bifunctional purine biosynthesis protein PurH | NIL-Wounding |
| Biosynthetic peptidoglycan transglycosylase | NIL-Wounding |
| C4-dicarboxylic acid transporter DauA | NIL-Wounding |
| Cadmium-transporting ATPase | NIL-Wounding |
| Carboxypeptidase | NIL-Wounding |
| Choline oxidase | NIL-Wounding |
| Chromosome partition protein Smc | NIL-Wounding |
| Cystathionine gamma-synthase | NIL-Wounding |
| D-3-phosphoglycerate dehydrogenase | NIL-Wounding |
| D-xylose-proton symporter | NIL-Wounding |
| Dihydropteroate synthase | NIL-Wounding |
| DNA ligase A | NIL-Wounding |
| DNA polymerase III subunit alpha | NIL-Wounding |
| DNA repair protein RadA | NIL-Wounding |
| DNA-directed RNA polymerase subunit beta EGKEAPGB_21235 | NIL-Wounding |
| Elongation factor G | NIL-Wounding |
| Energy-dependent translational throttle protein EttA | NIL-Wounding |
| Extracellular serine protease | NIL-Wounding |
| Fe(3+) dicitrate transport protein FecA | NIL-Wounding |
| Fimbrial subunit type 1 | NIL-Wounding |
| Formate dehydrogenase-O major subunit | NIL-Wounding |
| Fosfomycin resistance protein AbaF | NIL-Wounding |
| Fructose import ATP-binding protein FruK | NIL-Wounding |
| Fumarate hydratase class I, aerobic | NIL-Wounding |
| GABA permease | NIL-Wounding |
| Galactarate dehydratase (L-threo-forming) | NIL-Wounding |
| Gentisate transporter | NIL-Wounding |
| Glucans biosynthesis glucosyltransferase H | NIL-Wounding |
| Glucans biosynthesis protein G | NIL-Wounding |
| Glucosamine kinase | NIL-Wounding |
| Glutamine--fructose-6-phosphate aminotransferase [isomerizing] | NIL-Wounding |
| Glutamyl-tRNA(Gln) amidotransferase subunit A | NIL-Wounding |
| Glutathione import ATP-binding protein GsiA | NIL-Wounding |
| Glutathione-binding protein GsiB | NIL-Wounding |
| Glycine betaine transporter OpuD | NIL-Wounding |
| Guanine/hypoxanthine permease PbuG | NIL-Wounding |
| Haloalkane dehalogenase | NIL-Wounding |
| High-affinity choline transport protein | NIL-Wounding |
| High-affinity proline transporter PutP | NIL-Wounding |

| | |
|---|---|
| Histidinol-phosphate aminotransferase | NIL-Wounding |
| HTH-type transcriptional regulator KipR | NIL-Wounding |
| HTH-type transcriptional regulator MalT | NIL-Wounding |
| HTH-type transcriptional regulator SgrR | NIL-Wounding |
| Hydroxyacylglutathione hydrolase | NIL-Wounding |
| hypothetical protein | NIL-Wounding |
| Inosine-5-monophosphate dehydrogenase FGGNMKLN_18635 | NIL-Wounding |
| Inositol 2-dehydrogenase/D-chiro-inositol 3-dehydrogenase | NIL-Wounding |
| ISL3 family transposase ISPfr6 | NIL-Wounding |
| L-arabinose isomerase | NIL-Wounding |
| L-asparagine permease 2 | NIL-Wounding |
| L-Rhamnulokinase | NIL-Wounding |
| Lactose operon repressor | NIL-Wounding |
| Levanbiose-producing levanase | NIL-Wounding |
| Lipid A export ATP-binding/permease protein MsbA | NIL-Wounding |
| lipid II flippase MurJ | NIL-Wounding |
| Long-chain-fatty-acid--CoA ligase | NIL-Wounding |
| Lysine--tRNA ligase 1 | NIL-Wounding |
| Lysine-specific permease | NIL-Wounding |
| Magnesium-transporting ATPase, P-type 1 | NIL-Wounding |
| Major myo-inositol transporter IolT | NIL-Wounding |
| Malate synthase A | NIL-Wounding |
| Malate synthase G | NIL-Wounding |
| NAD(P)H dehydrogenase (quinone) | NIL-Wounding |
| NAD(P)H-quinone oxidoreductase subunit 2, chloroplastic | NIL-Wounding |
| Neutral endopeptidase | NIL-Wounding |
| Nitrilotriacetate monooxygenase component A | NIL-Wounding |
| Nitrite reductase [NAD(P)H] | NIL-Wounding |
| Oligo-1,6-glucosidase | NIL-Wounding |
| Outer membrane protein assembly factor BamA | NIL-Wounding |
| Peptidoglycan D,D-transpeptidase MrdA | NIL-Wounding |
| Phenylalanine--tRNA ligase beta subunit | NIL-Wounding |
| Phospho-2-dehydro-3-deoxyheptonate aldolase | NIL-Wounding |
| Phosphomethylpyrimidine synthase | NIL-Wounding |
| Polyisoprenyl-teichoic acid--peptidoglycan teichoic acid transferase TagU | NIL-Wounding |
| Polyphosphate kinase | NIL-Wounding |
| Polyprenol-phosphate-mannose-dependent alpha-(1-2)-phosphatidylinositol mannoside mannosyltransferase | NIL-Wounding |
| Proline--tRNA ligase | NIL-Wounding |
| Proline-specific permease ProY | NIL-Wounding |
| Protein translocase subunit SecD | NIL-Wounding |

| | |
|---|---|
| Protein UmuC | NIL-Wounding |
| Putative 3-oxopropanoate dehydrogenase | NIL-Wounding |
| putative AAA domain-containing protein | NIL-Wounding |
| putative ABC transporter ATP-binding protein YbiT | NIL-Wounding |
| putative ABC transporter ATP-binding protein YheS | NIL-Wounding |
| Putative acetyl-coenzyme A carboxylase carboxyl transferase subunit beta | NIL-Wounding |
| putative adenylyltransferase/sulfurtransferase MoeZ | NIL-Wounding |
| Putative amidase AmiB2 | NIL-Wounding |
| putative amino acid permease YhdG | NIL-Wounding |
| Putative cystathionine beta-synthase | NIL-Wounding |
| putative glycine dehydrogenase (decarboxylating) | NIL-Wounding |
| putative helicase HelY | NIL-Wounding |
| Putative multidrug export ATP-binding/permease protein | NIL-Wounding |
| putative oxidoreductase | NIL-Wounding |
| putative phosphomannomutase | NIL-Wounding |
| putative propionyl-CoA carboxylase beta chain 5 | NIL-Wounding |
| putative protein YhaP | NIL-Wounding |
| putative protein YihR | NIL-Wounding |
| putative siderophore transport system permease protein YfhA | NIL-Wounding |
| putative xanthine dehydrogenase subunit D | NIL-Wounding |
| putative zinc protease | NIL-Wounding |
| Putrescine oxidase | NIL-Wounding |
| Pyruvate carboxylase | NIL-Wounding |
| Sensor histidine kinase DcuS | NIL-Wounding |
| Sensor histidine kinase RcsC | NIL-Wounding |
| Sodium, potassium, lithium and rubidium/H(+) antiporter | NIL-Wounding |
| Swarming motility protein SwrC | NIL-Wounding |
| Thiol-disulfide oxidoreductase ResA | NIL-Wounding |
| Thioredoxin reductase | NIL-Wounding |
| Toluene efflux pump outer membrane protein TtgI | NIL-Wounding |
| Transcription-repair-coupling factor | NIL-Wounding |
| Transketolase | NIL-Wounding |
| Transketolase 1 | NIL-Wounding |
| Tryptophan 2-monooxygenase | NIL-Wounding |
| Uric acid transporter UacT | NIL-Wounding |
| Urocanate hydratase | NIL-Wounding |
| UvrABC system protein A | NIL-Wounding |
| Vitamin B12 import ATP-binding protein BtuD | NIL-Wounding |
| Vitamin B12 transporter BtuB | NIL-Wounding |
| Xylose import ATP-binding protein XylG | NIL-Wounding |
| Xylulose kinase | NIL-Wounding |

| | |
|---|---|
| Acyl carrier protein | NIL-Control |
| Biofilm dispersion protein BdlA | NIL-Control |
| Cytochrome c oxidase subunit 1 , bacteroid | NIL-Control |
| Error-prone DNA polymerase | NIL-Control |
| Ferredoxin--NADP reductase | NIL-Control |
| Ferric aerobactin receptor | NIL-Control |
| Glutathione-regulated potassium-efflux system protein KefC | NIL-Control |
| Glycine dehydrogenase (decarboxylating) | NIL-Control |
| Histidine--tRNA ligase | NIL-Control |
| hypothetical protein | NIL-Control |
| K(+)/H(+) antiporter NhaP | NIL-Control |
| L-2-hydroxyglutarate dehydrogenase | NIL-Control |
| Linear gramicidin synthase subunit B | NIL-Control |
| Lon protease | NIL-Control |
| Long-chain-fatty-acid--CoA ligase | NIL-Control |
| Metal-pseudopaline receptor CntO | NIL-Control |
| Phenylalanine--tRNA ligase beta subunit | NIL-Control |
| putative lipid II flippase MurJ | NIL-Control |
| putative peptidoglycan D,D-transpeptidase FtsI | NIL-Control |
| Sensor histidine kinase RcsC | NIL-Control |
| Type IV pilus biogenesis and competence protein PilQ | NIL-Control |
| Tyrocidine synthase 3 | NIL-Control |
| UDP-N-acetylmuramoyl-tripeptide--D-alanyl-D-alanine ligase | NIL-Control |

| **Gene (RNAi–Herbivory vs RNAi-Control)** | **Group with higher abundance** |
|---|---|
| 1,4-alpha-glucan branching enzyme GlgB | RNAi-Herbivory |
| 1,4-dihydroxy-2-naphthoyl-CoA synthase | RNAi-Herbivory |
| 2-methyl-1,2-propanediol dehydrogenase | RNAi-Herbivory |
| 3-[(3aS,4S,7aS)-7a-methyl-1,5-dioxo-octahydro-1H-inden-4-yl]propanoyl:CoA ligase | RNAi-Herbivory |
| 3-isopropylmalate dehydrogenase | RNAi-Herbivory |
| 3-ketoacyl-CoA thiolase FadI | RNAi-Herbivory |
| 3-ketosteroid-9-alpha-monooxygenase, oxygenase component | RNAi-Herbivory |
| 3-oxosteroid 1-dehydrogenase | RNAi-Herbivory |
| 3-succinoylsemialdehyde-pyridine dehydrogenase | RNAi-Herbivory |
| 6-deoxyerythronolide-B synthase EryA3, modules 5 and 6 | RNAi-Herbivory |
| 60 kDa chaperonin 1 | RNAi-Herbivory |
| ABC transporter ATP-binding/permease protein | RNAi-Herbivory |
| Aclacinomycin methylesterase RdmC | RNAi-Herbivory |
| Acyl-CoA dehydrogenase | RNAi-Herbivory |
| Aspartokinase | RNAi-Herbivory |

| | |
|---|---|
| ATP-dependent RNA helicase SrmB | RNAi-Herbivory |
| Baeyer-Villiger monooxygenase | RNAi-Herbivory |
| Bifunctional purine biosynthesis protein PurH | RNAi-Herbivory |
| Bifunctional uridylyltransferase/uridylyl-removing enzyme | RNAi-Herbivory |
| Cell division protein FtsQ | RNAi-Herbivory |
| Chaperone protein DnaK | RNAi-Herbivory |
| DNA gyrase subunit B | RNAi-Herbivory |
| DNA repair protein RecN | RNAi-Herbivory |
| DNA translocase FtsK | RNAi-Herbivory |
| Fatty acid ABC transporter ATP-binding/permease protein | RNAi-Herbivory |
| FO synthase | RNAi-Herbivory |
| Formate dehydrogenase H | RNAi-Herbivory |
| Glutamate synthase [NADPH] small chain | RNAi-Herbivory |
| Glutamate-1-semialdehyde 2,1-aminomutase | RNAi-Herbivory |
| hypothetical protein | RNAi-Herbivory |
| Inner membrane ABC transporter permease protein YdcV | RNAi-Herbivory |
| Iron import ATP-binding/permease protein IrtB | RNAi-Herbivory |
| Isoleucine--tRNA ligase | RNAi-Herbivory |
| Ketol-acid reductoisomerase (NADP(+)) | RNAi-Herbivory |
| L,D-transpeptidase 2 | RNAi-Herbivory |
| Limonene 1,2-monooxygenase | RNAi-Herbivory |
| Lipoprotein LprN | RNAi-Herbivory |
| Magnesium-transporting ATPase, P-type 1 | RNAi-Herbivory |
| Methylmalonyl-CoA carboxyltransferase 12S subunit | RNAi-Herbivory |
| Mycinamicin IV hydroxylase/epoxidase | RNAi-Herbivory |
| N-(2-amino-2-carboxyethyl)-L-glutamate synthase | RNAi-Herbivory |
| NADH-quinone oxidoreductase subunit G | RNAi-Herbivory |
| o-succinylbenzoate synthase | RNAi-Herbivory |
| Oxygen sensor histidine kinase response regulator DevS/DosS | RNAi-Herbivory |
| Phenolphthiocerol synthesis polyketide synthase type I Pks15/1 | RNAi-Herbivory |
| Phosphoribosylformylglycinamidine synthase subunit PurL | RNAi-Herbivory |
| Protein translocase subunit SecA 1 | RNAi-Herbivory |
| Protein translocase subunit SecF | RNAi-Herbivory |
| Putative acyl-CoA dehydrogenase FadE17 | RNAi-Herbivory |
| putative arabinosyltransferase C | RNAi-Herbivory |
| putative cytochrome c oxidase subunit 1 | RNAi-Herbivory |
| putative enoyl-CoA hydratase echA8 | RNAi-Herbivory |
| putative M18 family aminopeptidase 2 | RNAi-Herbivory |
| putative monooxygenase | RNAi-Herbivory |
| putative protein | RNAi-Herbivory |
| putative sensor histidine kinase TcrY | RNAi-Herbivory |

| | |
|---|---|
| Putative succinate-semialdehyde dehydrogenase [NADP(+)] 2 | RNAi-Herbivory |
| putative transporter | RNAi-Herbivory |
| Pyruvate dehydrogenase [ubiquinone] | RNAi-Herbivory |
| Respiratory nitrate reductase 2 beta chain | RNAi-Herbivory |
| Siderophore exporter MmpL4 | RNAi-Herbivory |
| Signal recognition particle protein | RNAi-Herbivory |
| Steroid C26-monooxygenase | RNAi-Herbivory |
| Succinyl-diaminopimelate desuccinylase | RNAi-Herbivory |
| Terminal beta-(1->2)-arabinofuranosyltransferase | RNAi-Herbivory |
| Urease subunit alpha | RNAi-Herbivory |
| 2-octaprenylphenol hydroxylase | RNAi-Control |
| 2,6-dihydropseudooxynicotine hydrolase | RNAi-Control |
| 4-cresol dehydrogenase [hydroxylating] flavoprotein subunit | RNAi-Control |
| 4-hydroxybenzoate transporter PcaK | RNAi-Control |
| 4-hydroxythreonine-4-phosphate dehydrogenase | RNAi-Control |
| 5-deoxy-glucuronate isomerase | RNAi-Control |
| 8-amino-7-oxononanoate synthase | RNAi-Control |
| Acetolactate synthase isozyme 1 large subunit | RNAi-Control |
| Acyl-CoA dehydrogenase | RNAi-Control |
| Adaptive-response sensory-kinase SasA | RNAi-Control |
| Aralkylamine dehydrogenase heavy chain | RNAi-Control |
| Aspartate kinase | RNAi-Control |
| Beta-barrel assembly-enhancing protease | RNAi-Control |
| Beta-lactamase | RNAi-Control |
| Beta-xylosidase | RNAi-Control |
| Chaperone SurA | RNAi-Control |
| Coniferyl aldehyde dehydrogenase | RNAi-Control |
| D-amino acid dehydrogenase | RNAi-Control |
| D-malate dehydrogenase [decarboxylating] | RNAi-Control |
| D-tagatose-1,6-bisphosphate aldolase subunit KbaZ | RNAi-Control |
| DNA gyrase subunit B | RNAi-Control |
| DNA topoisomerase 3 | RNAi-Control |
| Endonuclease MutS2 | RNAi-Control |
| Flavin-dependent tryptophan halogenase RebH | RNAi-Control |
| Galactose-proton symporter | RNAi-Control |
| GDP-L-fucose synthase | RNAi-Control |
| Glutamate-pyruvate aminotransferase AlaA | RNAi-Control |
| Glycine betaine-binding protein YehZ | RNAi-Control |
| Hercynine oxygenase | RNAi-Control |
| Histidinol-phosphate aminotransferase | RNAi-Control |
| hypothetical protein | RNAi-Control |

| | |
|---|---|
| Inner membrane protein YedI | RNAi-Control |
| Inner membrane transport protein YdhP | RNAi-Control |
| IS30 family transposase IS1088 | RNAi-Control |
| L-cystine uptake protein TcyP | RNAi-Control |
| Lon protease 2 | RNAi-Control |
| Maleylacetate reductase | RNAi-Control |
| Mannuronan C5-epimerase AlgE2 | RNAi-Control |
| Miniconductance mechanosensitive channel YbdG | RNAi-Control |
| Multidrug resistance protein MdtA | RNAi-Control |
| N-substituted formamide deformylase | RNAi-Control |
| NAD-dependent protein deacylase | RNAi-Control |
| Nicotinamide-nucleotide amidohydrolase PncC | RNAi-Control |
| Oligopeptide-binding protein AppA | RNAi-Control |
| PCP degradation transcriptional activation protein | RNAi-Control |
| Phosphate-binding protein PstS | RNAi-Control |
| Polyamine aminopropyltransferase | RNAi-Control |
| Protein GltF | RNAi-Control |
| Putative acyl-CoA dehydrogenase AidB | RNAi-Control |
| Putative aldehyde dehydrogenase AldA | RNAi-Control |
| putative diacyglycerol O-acyltransferase tgs1 | RNAi-Control |
| Putative glycoside/cation symporter YagG | RNAi-Control |
| Putative metabolite transport protein YjhB | RNAi-Control |
| putative signaling protein | RNAi-Control |
| Ribose operon repressor | RNAi-Control |
| RNA polymerase-associated protein RapA | RNAi-Control |
| Sensor histidine kinase GlrK | RNAi-Control |
| Single-stranded-DNA-specific exonuclease RecJ | RNAi-Control |
| Thiosulfate sulfurtransferase YnjE | RNAi-Control |
| Tyrosine recombinase XerC | RNAi-Control |
| Vitamin B12 transporter BtuB | RNAi-Control |
| Xylulose kinase | RNAi-Control |
| Zinc transport protein ZntB | RNAi-Control |

| **Gene (RNAi-Wounding vs RNAi-Control)** | **Group with higher abundance** |
|---|---|
| 4-aminobutyrate aminotransferase | RNAi-Wounding |
| Adaptive-response sensory-kinase SasA | RNAi-Wounding |
| ATP-dependent Clp protease ATP-binding subunit ClpC1 | RNAi-Wounding |
| HTH-type transcriptional activator TipA | RNAi-Wounding |
| hypothetical protein | RNAi-Wounding |
| putative deoxyribonuclease RhsA | RNAi-Wounding |

| | |
|---|---|
| Secretin GspD 2 | RNAi-Wounding |
| hypothetical protein | RNAi-Control |
| IS3 family transposase ISMsm7 | RNAi-Control |

**Methods S1 Plant material and growth conditions.**

*T. koksaghyz* was germinated in propagation substrate (VM, Einheitserde, https://www.einheitserde.de/) and subsequently transferred to 3 L pots filled with a 1:1 mixture of standard soil (ED73, Einheitserde) and sand, except for the microbiome experiment for which a 4:1 mixture of natural field soil and sand was used. During cultivation and until five days prior to experiments, organic farming non-systemic insecticides (Spruzit Neudorff, Naturalis® and Para Sommer Dr. Stähler) were applied above-ground to control fungus gnat, white fly, spider mite and thrips infestation. *Daucus carota* seeds were germinated and cultivated in the propagation substrate.

As *T. koksaghyz* is self-incompatible, seed material used in our experiments was obtained by hand pollinating the T1 generation of two independent *Tk*CPTL1-RNAi lines (RNAi-A and RNAi-B) with *T. koksaghyz* wild type pollen. For details on identification of RNAi and near-isogenic line (NIL) plants see Supporting Information Methods S2.

**Methods S2 Identification of transgenic rubber-depleted plants**

Transgenic rubber-depleted *Tk*CPTL1-RNAi and NIL plants were identified based on PCR amplification of two *Tk*CPTL1-RNAi vector fragments using gDNA of 3-week-old T2 leaf material (KAPA 3G plant kit; Roche, Basel, Switzerland) using vector specific primers pREF-seq-fwd and P1 and RNAi fragment primers flanking both sides of the construct. Plants giving none or one band were assigned as NIL plants. As a technical control, the housekeeping gene glycerine aldehyde-3-phosphate dehydrogenase (GAPDH) was amplified. PCR products were run on a 1% agarose gel. Corresponding plants of PCR products showing both fragments (as well as GAPDH band) were assigned as rubber-depleted RNAi plants. The promoter used is laticifer-predominant (pREF = promoter of the rubber elongation factor gene).

pREF-seq-fwd (CGTAGCCAAGCGATATTCAATATC) + *Tk*CPTL1-RNAi-NcoI-fwd AAACCATGGCTTATAATAAATGAAATTGT): 575 bp

*Tk*CPTL1-RNAi-NcoI-fwd + P1 (ATCATGCGATCATAGGCGTC): 481 bp

GAPDH_fwd (CTTCAGAGAGATGATGTT) + GAPDH_rev (CTTCCACCTCTCCAGTCCTT)

**Methods S3 Isolation of pure *cis*-1,4-polyisoprene from *T. koksaghyz* crude rubber nuggets and preparation of *cis*-1,4-polyisoprene solution for chemical supplementation experiments with carrots seedlings.**

In order to obtain phospholipid- and triterpene-free *cis*-1,4-polyisoprene used for supplementation experiments, 22 g of crude rubber nuggets from *T. koksaghyz* wildtype, see also (Kreuzberger *et al.*, 2016; Eggert *et al.*, 2018), were placed into MeOH/Chloroform (2:1). Extraction solution was replaced every 48 h for 3 consecutive times resulting in 18 g of pure *cis*-1,4-polyisoprene (based on (Bonfils *et al.*, 2007; Bergmann *et al.*, 2018). The resulting nugget was oven dried for 24 h at 40 °C. We then generated a 1% (w/v) *cis*-1,4,-polyisoprene/chloroform slurry used for supplementation experiments.

**Methods S4 Chemical supplementation and genetic modification of triterpenes within the latex fraction of *T. koksaghyz*.**

***Identification of transgenic triterpene-reduced plants***. Transgenic triterpene-reduced *TkOSC*-RNAi and NIL plants were identified based on PCR amplification of *Tk*OSC-RNAi vector fragments using gDNA of 3-week-old T1 leaf material (KAPA 3G plant kit; Roche, Basel, Switzerland) using vector specific primers P2 and P3 and RNAi fragment primers flanking both sides of the construct. Plants giving none or one band were assigned as NIL plants. The promoter used is laticifer-predominant (pREF = promoter of the rubber elongation factor gene).

*TkOSC*1-RNAi:

P2 (TACCTTCCCACAATTCGTCG) + *TkOSC1*-RNAi-rev-XhoI (AAACTCGAGATCCCAAGCTTGAATCGCAC): 433bp

P3 (CAGGTATTGGATCCTAGGTG) + *TkOSC1*-RNAi-rev-XhoI (AAACTCGAGATCCCAAGCTTGAATCGCAC): 300bp

*TkOSC*-RNAi (OSC unspecific):

P2 (TACCTTCCCACAATTCGTCG) + *TkOSC1*-RNAi-rev-XhoI (AAACTCGAGTCTCCAAGCGCCCATAGCGG): 753bp

P3 (CAGGTATTGGATCCTAGGTG) + *TkOSC1*-RNAi-rev-XhoI (AAACTCGAGTCTCCAAGCGCCCATAGCGG): 600bp

***Plant performance experiment.*** We first tested whether triterpenes within the latex fraction affect plant performance under herbivory *in planta* using two independent triterpene-reduced RNAi lines (L2 and L3, (van Deenen *et al.*, 2019)), analogous to the experiment using the rubber-depleted RNAi lines. Here, plants were precultivated as described before. In each plant line, half of RNAi and NIL plants were infested with a starved and pre-weighed larva, originating from Germany (n=10 plants per treatment per plant line) for nine days. One herbivory replicate per genotype for L3 was lost due to dying larvae. Above and below ground fresh weight of plants was analyzed using linear models with the formula '*biomass~plant_genotype*herbivory_treatment'*.

Pairwise comparisons between treatments within each plant genotype were adjusted using the FDR method with the package emmeans (Lenth, 2022) using the formula 'pairwise~herbivory_treatment|plant_genotype'. Larval weight gain of larvae was analyzed using a Wilcoxon test. As intrinsic biomass of the two plant lines differed among each other, datasets were analyzed separately using the *lme4* package (Bates *et al.*, 2015).

***Larval performance experiment ex planta.*** In order to test whether a *T. koksaghyz* latex triterpene, lupeol (Unland *et al.*, 2018; Pütter *et al.*, 2019), affects larval diet consumption and weight gain, we allowed *M. melolontha* to feed on artificial diet supplemented with lupeol or the solvent control. Lupeol increases two-fold based on root dry weight in the natural rubber-depleted *Tk*CPTL1 (Niephaus *et al.*, 2019). Larvae, originating from Germany, were starved for 72 h before being allowed to feed for 48 h on 400 mg semiartificial diet cubes (recipe see prior experiment) supplemented with an ecologically relevant amount of lupeol (Extrasynthese, France), 0.1% per fresh cube, or solvent only (hexane) (n=20 each). Solvent was allowed to evaporate prior to feeding. Diet cubes were each placed into a 250 ml plastic beaker with one larva each. Larvae not feeding at all were excluded from the data set resulting in n=17 control and n=19 lupeol. Diet consumption and larval weight gain were analyzed with Wilcoxon signed-rank tests.

**Methods S5 Origin and storage of soil used for microbiome experiment**

Soil was provided by Fred Eickmeyer (ESKUSA, Parkstetten, Germany). Soil originated from the field “Stahl Aufeld” near Parkstetten (Germany). Here, *T. koksaghyz* had been cultivated for several years. Soil was collected after dandelion plants had been harvested in early autumn (medium heavy soil, using the upper 30 cm soil after plowing) and soil was sent to us in late November 2018, where it was stored in buckets at ambient temperature in the dark until used for *T. koksaghyz* cultivation in our glasshouse starting at the end of January 2019. Soil was sieved and homogenized prior to usage. Once seedlings were repotted to 3-liter pots, homogenized field soil was mixed with 20% sand.

**Methods S6 Handling of samples for microbiome study and statistical analysis of biomass data**

Two days prior to harvest, plants were not watered anymore to facilitate harvest. Plants were harvested 14 days after infestation and larval weight was determined. Roots were slightly shaken to remove loose soil. Then, root and shoot fresh weights were determined. To obtain the rhizosphere fraction, roots were washed in three consecutive steps with 30 ml sterile water similar to (Hu *et al.*, 2018). The washing fractions (I, II, III) and washed roots were immediately frozen in liquid nitrogen, stored at – 20 °C and subsequently freeze-dried. Dried material of washing fractions I, II, III of each plant were combined into one sample and defined as the rhizosphere sample. To obtain the root microbiome fraction (endo- and ectophytes), lyophilized roots were ground using an analysis mill (IKA A11 basic A11 BS00, Staufen im Breisgau, Germany; 2 rounds for 15 sec each); the grinder was cleaned with air flow in between samples and thoroughly washed with ultrapure water in between treatments. All samples were stored at room temperature in the dark until further processing. For statistical analysis of plant and larval performance under herbivory and control treatment, we excluded spontaneously flowering and heavily wilted plants (NIL-A non-infested n=9, NIL-A infested n=8, RNAi-A non-infested n=9, RNAi-A infested n=8). For the subset of n=6 used for the microbiome analysis, we also analyzed shoot and root biomass accumulation including herbivory and control treatment using linear models with the formula '*biomass~plant_genotype*treatment'.* Pairwise comparisons between treatments within each plant genotype were adjusted using the FDR method with the package emmeans (Lenth, 2022) using the formula 'pairwise~treatment|plant_genotype'. The effect of natural rubber silencing on larval weight gain was analyzed with a Wilcoxon signed-rank test.

**Notes S1 Taxa analysis**

Given the differences in the microbial community structure in response to plant genotype and treatment, we tested which microbial taxa differ between each treatment and the respective control group using two different methods. As a first test, we contrasted each treatment (herbivory or wounding) against the control group within each plant genotype using DESeq2. In the rhizosphere bacterial community, the number of OTUs that varied between treatment and control was different from random for all genotype * treatment combinations ($P$<0.001, Supporting Information Table S6). When contrasting NIL plants exposed to herbivory towards their control, we did not find any differentially abundant OTUs. However, when contrasting wounded NIL plants towards the control we found 2 OTUs more abundant in wounded plants (one *Pseudonocardia* and one unidentified) and 115 OTUs more abundant in control plants (Supporting Information Table S7). When focusing on RNAi plants, 5 OTUs were found more abundant in plants exposed to herbivory than their control (*Alicyclobacillus*, *Devosia*, *Emticicia*, *Microbacterium*, *Nocardioldes*), while 2 OTUs were more abundant in the control group (*Pseudomonas* and one unidentified). Similarly, 1 OTUs identified as *Pseudomonas* was more abundant in control plants compared to wounded RNAi plants (Supporting Information Table S7). When focusing on the rhizosphere fungal community, this approach identified only 4 OTUs that were more abundant in wounded NIL plants compared to their control (*Bjerkandera*, *Gallinipes*, *Toxicocladosporium*, and one unidentified) while no differentially abundant OTUs were identified for the other groups (Supporting Information Table S8). When focusing on root samples, only one bacterial OTU was identified to be more abundant in RNAi plants exposed to herbivory than control plants (*Arthrobacter*), while no fungal OTUs were found to be differentially abundant (Supporting Information Table S9-S10).

As a second test, we aggregated our dataset at the genus level for each sample, and we focused on those that on average represent more than 1% of the community across all samples. For each plant genotype * treatment combination, we then built a linear model to test differences in the relative abundance of each microbial genus between treated plants (herbivory or wounding) and their control. When focusing on rhizosphere samples, we found an increase in relative abundance of *Arthrobacter* in NIL plants exposed to both herbivory and wounding, while in wounded plants the relative abundance of *Rhizobium* decreased. In the rhizosphere of RNAi plants, we observed an increase in *Mesorhizobium* and *Mycobacterium* when plants were exposed to herbivory, and an increase of *Microbacterium*, *Mycobacterium*, *Novosphingobium* and *Pseudomonas* when plants were exposed to wounding (Supporting Information Table S11). When focusing on the bacterial community of plant roots, in NIL plants we observed and increase in the relative abundance of *Bacillus* and Mycobacterium when plants were exposed to herbivory, while when plants were exposed to wounding, we observed a decrease in abundance of *Cellvibrio*, *Novosphingobium*, *Rhizobium* and *Sphingobium*. In RNAi plants, we observed a decrease in abundance of *Bacillus* and *Paenibacillus* when plants were exposed to both herbivory and wounding, and a decrease in the abundance of *Pseudomonas* when plants were exposed to wounding (Supporting Information Table S12).
